# Dynamics of gene expression and chromatin marking during cell state transition

**DOI:** 10.1101/2020.11.20.391524

**Authors:** Beatrice Borsari, Amaya Abad, Cecilia C. Klein, Ramil Nurtdinov, Alexandre Esteban, Emilio Palumbo, Marina Ruiz-Romero, María Sanz, Bruna R. Correa, Rory Johnson, Sílvia Pérez-Lluch, Roderic Guigó

## Abstract

We have monitored the transcriptomic and epigenomic status of cells at twelve time-points during the transdifferentiation of human pre-B cells into macrophages. Using this data, we have investigated some fundamental questions regarding the role of chromatin in gene expression. We have found that, over time, genes are characterized by a limited number of chromatin states (combinations of histone modifications), and that, consistently, chromatin changes over genes tend to occur in a coordinated manner. We have observed strong association between these changes and gene expression only at the time of initial gene activation. Activation is preceded by H3K4me1 and H3K4me2, and followed in a precise order by most other histone modifications. Further changes in gene expression, comparable or even stronger than those at initial activation, occur without associated changes in histone modifications. The data generated here constitutes, thus, a unique resource to investigate transcriptomic and epigenomic dynamics during a differentiation process.

## Introduction

Chromatin is the complex of DNA, histone and non-histone proteins that constitutes the chromosomes found in the nucleus of eukaryotic cells. Post-translational modifications (PTMs) of histone proteins, together with other epigenetic features, can alter the overall chromatin structure and are thought to play a critical role in the regulation of all DNA-based processes. In particular, interest has grown in understanding the relationship between chromatin and transcriptional regulation.

Several histone marks have been associated with either active or silent gene expression. For instance, high levels of H3K27ac and H3K4me1 are considered a feature of active transcriptional enhancers (Hon et al., 2009), whereas active promoters are typically marked by H3K4me3 (Barski et al., 2007; Schneider et al., 2004). Conversely, features of constitutive and facultative heterochromatin correlate with levels of H3K9me3 and H3K27me3, respectively (Trojer and Reinberg, 2007; Hansen et al., 2008). This is strongly suggestive of an association between gene expression and chromatin modifications. According to the histone code hypothesis (Strahl and Allis, 2000), distinct combinations of histone modifications over regulatory regions — associated with specific arrangement of transcription factors — confer to each gene a unique temporal and spatial transcriptional program. Based on this hypothesis, methods to predict gene expression from combinations of different histone marks have been developed with great accuracy, even when the predictions are obtained in a cell type other than the one in which the model is inferred (Karlić et al., 2010; Dong et al., 2012).

The majority of these predictions are conducted in steady-state conditions, and therefore do not track the association between gene expression and histone marks over time. Studies along time, however, are essential to decipher the mechanisms behind transcriptional control and maintenance, since an appropriate balance of stability and dynamics in epigenetic features seems to be required for accurate gene expression (Greer and Shi, 2012). Interestingly, a number of studies in different species and biological models have highlighted a degree of correlation between gene expression and chromatin marks over time substantially lower than what previously described in steady-state conditions. For instance, during fruit fly development, around 34% of the expressed genes lack H3K4me3 at their promoters (Nègre et al., 2011), while transcription can occur in the absence of most active marks (Hödl and Basler, 2012; Pérez-Lluch et al., 2015). It has also been reported that, upon stimulation, changes in gene expression are not always accompanied by changes in histone modifications (Vandenbon et al., 2018), and that chromatin marks do not represent linear measures of transcriptional activity (Le Martelot et al., 2012; Wang et al., 2019). Overall, it has been suggested that the contribution of chromatin to gene expression depends largely on the promoter architecture of genes (Rach et al., 2011).

Time-series studies have also striven to elucidate the temporal ordering in which transcription factor (TF) binding, deposition of histone marks and RNA Polymerase recruitment occur at both enhancer and promoter regions. For instance, it has been reported that enhancers required for hematopoietic differentiation are already primed with H3K4me1 in multipotent progenitors (Mercer et al., 2011). However, *de novo* enhancers’ transcription seems to precede local deposition of H3K4me1 and H3K4me2 marks (Kaikkonen et al., 2013). Furthermore, deposition of H3K4me1 is dispensable for either enhancer or promoter transcription, and does not affect the maintenance of transcriptional programs (Dorighi et al., 2017; Cao et al., 2018).

Nevertheless, most time-series studies so far have monitored a few histone modifications in a limited number of time-points. To address these limitations, here we have generated gene expression profiles and maps of nine histone modifications at twelve time-points along a controlled cellular differentiation process: the induced transdifferentiation of human BLaER1 cells into macrophages (Rapino et al., 2013). BLaER1 is a human B-cell precursor leukemia cell line, stably transfected with a construct containing cEBPα fused with the estrogen hormone receptor binding domain (Rapino et al., 2013). These cells are able to transdifferentiate into functional macrophages at a high efficiency rate upon induction with beta-estradiol, which induces the internalization of the transcription factor into the nucleus, promoting massive transcriptomic changes. We believe that the data that we have generated constitutes an unprecedented resource in the field of time-series transcriptional and chromatin studies.

Analysis of these data reveals that the large steady-state associations between gene expression and chromatin marking previously reported are partially artifactual, and mainly arise from the constrained nature of the transcriptome and the epigenome. We found that only a limited number of combinations of histone modifications are actually marking the genes, defining the major genic chromatin states in the human genome. Genes tend to remain in the same state throughout the entire transdifferentiation process, even those that change expression substantially. We have also observed substantial chromatin changes that are not necessarily accompanied by changes in gene expression, suggesting that epigenetic modifications contribute to cell state in a manner that cannot be fully recapituted by gene expression. We did find, however, a strong association between chromatin marking and expression at the time of initial gene activation. We have been able to determine the precise order of histone modifications at that time, and found that only H3K4me1 and H3K4me2 appear to be deposited prior to gene activation. Further changes in gene expression, comparable or even stronger than those at gene activation, seem to be mostly uncoupled from changes in histone modifications.

## Results

### A rich resource for time-series analysis of chromatin and gene expression dynamics

To investigate the temporal interplay between transcriptional activity and chromatin marking during the transdifferentiation of BlaER1 cells into macrophages (Rapino et al., 2013), we monitored this process at 12 time-points, from 0 to 168 hours post-induction (p.i.) (Fig. 1a). Reciprocal regulation of B-cell and macrophage antigens CD19 and Mac-1, respectively, was assessed by flow cytometry throughout the process (Fig. S1a). For each time-point we characterized, in two biological replicates, the whole cell RNA-seq gene expression profiles and the ChIP-seq maps of nine histone post-translational modifications. Besides the six marks (H3K4me1, H3K4me3, H3K27ac, H3K27me3, H3K36me3 and H3K9me3) endorsed by the reference epigenome criteria (International Human Epigenome Consortium, http://ihec-epigenomes.org/research/reference-epigenome-standards/), we have profiled H3K4me2, H3K9ac and H4K20me1 (Fig. 1b). This has allowed us to characterize the interchange between different degrees of lysine four methylation over time, but also to compare acetylation patterns on distinct lysine residues, and to explore the alternation of broad marks over actively transcribed gene bodies. In addition, we have generated, for each time-point, ChIP-seq profiles of the transcription factor cEBPα, RNA-seq data from the cytosol and the nucleus, as well as riboprofiling and proteomics maps (Correa *et al*., in preparation).

**Figure 1:**
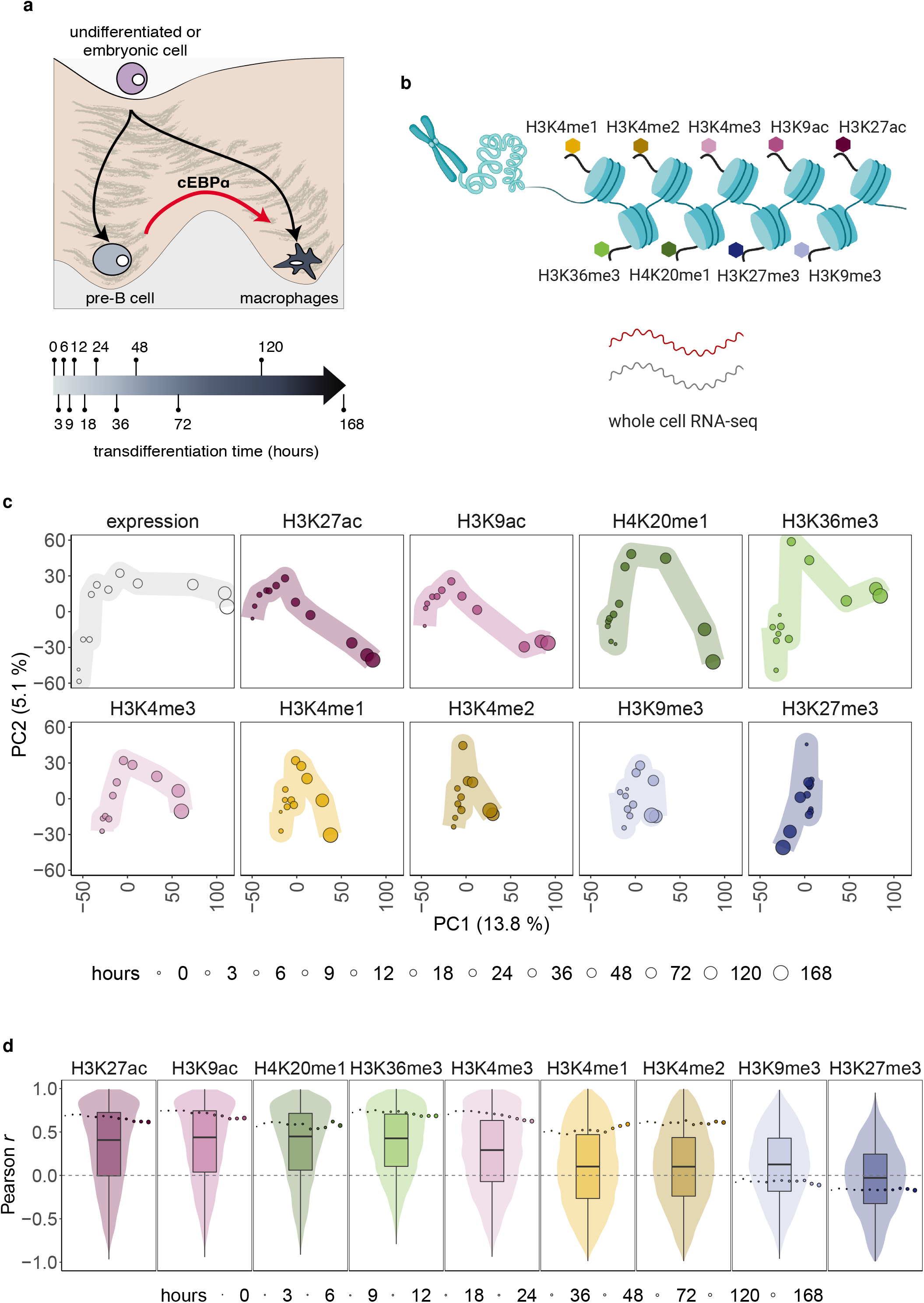
Global behaviour and relationship between chromatin and expression during transdifferentiation —. See also Figs. S1-2-6, Tables S1-2. **a**: The transdifferentiation of human pre-B cells into macrophages lasts a period of seven days, which we monitored at twelve time-points. **b**: We have performed ChIP-seq of nine histone modifications and RNA-seq in whole-cell fraction, at twelve time-points along the process of transdifferentiation. All experiments were performed in two biological replicates. **c**: Trajectories of transdifferentiation derived from a Principal Component Analysis performed jointly on timeseries gene expression and chromatin marks’ profiles. **d**: Correlations between levels of gene expression and histone marks. For a given mark and for each of the twelve time-points, we computed the steady-state Pearson *r* value between the vector of expression levels and the vector of chromatin signals corresponding to the 12,248 genes. These twelve correlation values are represented by single dots, the size of the dot being proportional to the hours of the corresponding time-point. The median Pearson r values for each mark are: H3K27ac: 0.67; H3K9ac: 0.72; H4K20me1: 0.59; H3K36me3: 0.72; H3K4me3: 0.70; H3K4me1: 0.51; H3K4me2: 0.61; H3K9me3: −0.07; H3K27me3: −0.17. In the case of time-course correlations, we obtained a Pearson *r* value for each expressed gene, and the distributions for all genes are represented by violin and box plots. Median Pearson *r* values across genes for each mark are: H3K27ac: 0.41; H3K9ac: 0.44; H4K20me1: 0.45; H3K36me3: 0.43; H3K4me3: 0.29; H3K4me1: 0.10; H3K4me2: 0.10; H3K9me3: 0.13; H3K27me3: −0.03.

To avoid any bias due to differences in the transdifferentiation process between experiments, a crucial component of our experimental design is that the RNA and the chromatin to perform immunoprecipitations with all histone marks were obtained from the same pool of cells in each biological replicate (see Methods). To efficiently and reproducibly analyze the wealth of data generated in a controlled environment, we developed *ChIP-nf* (https://github.com/guigolab/chip-nf), a pipeline implemented in NextFlow (DI Tommaso et al., 2017, see Methods).

### Gene expression recapitulates transdifferentiation more accurately than chromatin

To characterize gene expression and histone modifications’ profiles during the pre-B cell transdifferentiation process, we selected 12,248 genes — out of 19,831 protein-coding genes annotated in Gencode (Frankish et al., 2019) version 24 — that were either expressed in at least one time-point (≥ 5 TPM, 10,696 genes), or silent all along the process (0 TPM in all time-points, 1,552 genes) (Fig. S1b). Within expressed genes, we identified 8,030 genes characterized by significant changes in their expression profiles over time (differentially expressed, DE; Fig. S1b; see Methods). Half of these genes are down-regulated during the process, 25% are up-regulated, and for the remaining 25% we observed transient increases (peaking) and decreases (bending) in expression. 2,666 expressed genes do not display changes in expression over time (stably expressed).

For every gene in these sets, we also computed the level of each histone modification at a specific time-point, either over the gene body in the case of H3K36me3 and H4K20me1, or at promoter regions (± 2 Kb with respect to the transcription start site) for the remaining marks (Fig. S1c, see Methods). Roughly all expressed genes are marked by the canonical active histone modifications, whereas the proportion of silent genes showing peaks of these marks is low, except for H3K4me1 and H3K4me2 (Table S1). Unexpectedly, marks typically associated with silent transcription (H3K9me3 and H3K27me3) are not abundant in either expressed or silent genes.

To visually summarize the gene expression and individual histone modification profiles during transdifferentiation, we performed Principal Component Analysis (PCA), in which we plotted the 12 time-points based on these profiles (Fig. 1c). Even though the PCA was performed jointly on gene expression and all chromatin marks — which show different patterns of variation —, the first two principal components (PC1 and PC2) still capture about one fifth of the total variance of the data. Whereas gene expression is able to recapitulate the process in the space of the first two principal components, the chromatin marks are less resolutive, with H3K27ac, H3K9ac and H4K20me1 showing the clearest trends. The trajectory of gene expression in the PCA space suggests that the process occurs in two different transcriptional phases, with PC1 explaining the main differences between pre-B cells and macrophages, and PC2 representing early transcriptional changes within the first 24 hours of transdifferentiation. Instead, for several chromatin marks we observed parabolic trajectories, with PC2 mainly separating the intermediate stages of transdifferentiation from the differentiated cell types. Genes contributing to PC1 are mostly up- or down-regulated (Fig. S1d), and display significant enrichment in Gene Ontology terms associated with immune response and cell motility (Table S2). Instead, PC2-contributing genes perform functions related to nucleic acids metabolism and protein modification (Table S2), and comprise a large proportion of genes either displaying no changes in gene expression, or presenting transient increases or decreases (Fig. S1d). Taken all together, these results suggest that while there are major changes in gene expression and chromatin leading from one differentiated cell type to another (PC1), there are also changes that may be involved in a transient de-differentiation from pre-B cells into an intermediate state, and in the re-differentiation into macrophages (PC2), with a distinct contribution of expression and chromatin marks.

### The association between chromatin marking and gene expression is overestimated by correlations computed in steady-state conditions

We computed, at each time-point, the steady-state correlation between levels of expression and histone modifications across the set of 12,248 genes (Fig. 1d). As previously observed, we found a strong positive correlation for most active marks (median Pearson *r* value across time-points between 0.51 and 0.72), and a (weak) negative correlation for the repressive marks H3K9me3 and H3K27me3 (−0.07 and −0.17, respectively). However, when computing, for individual genes, the correlation between expression and chromatin profiles through time (time-course correlations), the values are substantially lower for active marks (median Pearson *r* ranging between 0.10 and 0.45), and higher for repressive marks (0.13 and −0.03 for H3K9me3 and H3K27me3, respectively; Fig. 1d). Remarkably, for H3K9me3 the time-course correlation with expression is positive, in contrast to what has been previously described (Ninova et al., 2019), and that we also measured in steady states.

It appears, therefore, that correlations measured in steady-state conditions artificially inflate the true degree of association between gene expression and chromatin modifications. This can be dramatically seen by randomizing the real temporal association between gene expression and chromatin marks. Within each gene’s time-series profile, we permuted histone modification levels among time-points, while keeping the actual gene expression values (see Methods; for an example with H3K4me3, compare upper and lower panels in Fig. S2a). As expected, the average time-course correlation is zero for all marks (Fig. S2b). However, the steady-state correlations are unexpectedly large for canonically active marks upon randomization, despite the fact that any meaningful association between gene expression and chromatin marks has been eliminated (Figs. S2a lower panel, S2b). This is likely due to a considerable fraction of genes displaying stable expression and chromatin profiles over time, which are either relatively highly expressed and marked (housekeeping genes, Pervouchine et al., 2015), or silent and not marked. Indeed, after removing the genes with silent or stable expression profiles over time, the steady-state correlations (Fig. S2c) are lower compared to those computed on the entire set of genes (Fig. 1d), and become more similar to the time-course correlations.

### Genes are characterized by a limited number of major chromatin states, which are more stable than expression

Next, we investigated the dynamics of chromatin marking during transdifferentiation. Towards that end, we summarized the chromatin state of each gene at each time-point, by building a multivariate Hidden Markov Model (HMM) on the signal of the nine histone marks along the twelve transdifferentiation points. More specifically, we produced a segmentation of the transdifferentiation time by assigning a given chromatin state to each gene at each time-point. This is in contrast to previous uses of HMMs in the field, where the segmentation is produced along the genome sequence by assigning a given chromatin state to every genome interval (Ernst and Kellis, 2012; Hoffman et al., 2012; Song and Chen, 2015; Zhang et al., 2016; Zhang and Hardison, 2017). We explored configurations with up to twenty different states, and found that five states are a good compromise between optimizing the likelihood of the model and the number of states capturing the epigenetic status of genes (Figs. S3a and 2a, see Methods). These five states correspond to the major combinations of histone modifications in which genes can be found (major chromatin states): a) absence of marking (with the exception, in some cases, of moderate marking by H3K9me3), b) low marking (mono and di-methylation of H3K4), c) bivalent marking (mostly marking by H3K4me1, H3K4me2 and/or H3K27me3), d) canonical active marking (all canonical active marks) and e) strong canonical active marking in the presence of H3K9me3 signal. These states (from a to e) correspond to increasing marking by canonically active histone modifications, with the exception of the bivalent marking state (c), which is also characterized by high H3K27me3 signal. These results suggest that only a limited number of combinations of marks can co-occur in a given gene at a given time-point. They also suggest that marking by H3K4me1 and H3K4me2 appears to be a precondition for marking by any other active histone modification, since for none of the configurations that we have explored, we have found states in which there is marking by an active histone modification without H3K4me1 and H3K4me2. The most frequent states among expressed genes are active and strong active marking (d and e, respectively), while the most frequent state among silent genes is absence of marking (a) (Fig. S3b).

Hierarchical clustering of genes based on the sequence of the five states along the twelve time-points revealed a limited number of temporal chromatin state profiles (Fig. 2b-c). Most of the genes remain in the same chromatin state during transdifferentiation (constant state profiles), irrespective of whether they are stably (79%) or differentially expressed (70%) during transdifferentiation (Fig. 2d, left panels). Thus, during the process, most changes in gene expression are not accompanied by chromatin changes.

**Figure 2:**
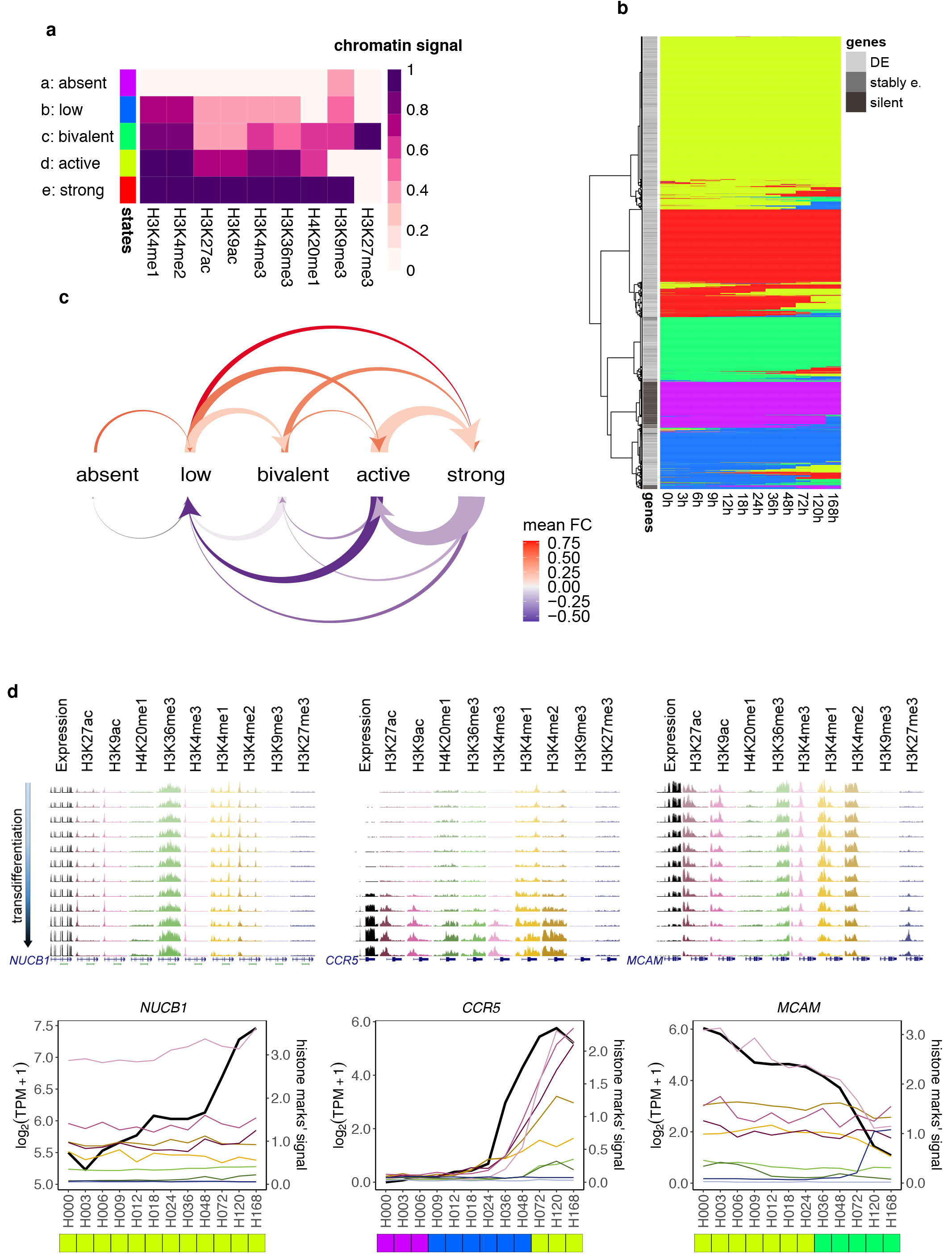
Genes are characterized by a limited number of major chromatin states, which are more stable than expression —. See also Fig. S3. **a**: A five-state multivariate HMM. Each state is defined by a combination of histone marks. We report the histone marks’ signals corresponding to each state. The states are sorted by increasing level of marking averaged over the nine histone modifications, with a and e states characterized by the lowest and highest average level of marking, respectively. **b**: Heatmap representing the hierarchical clustering of the HMM profiles built along the transdifferentiation process for the 12,248 genes. **c**: Arc diagram representing the types of state transitions observed in the HMM-sequence profiles of DE genes. The size of the arrow base is proportional to the number of genes reporting a given transition. Only transitions involving ≥ 10 genes are shown. We tested, for the sets of genes reporting each type of transition, the significance in gene expression fold-change (FC) (Wilcoxon Rank-Sum paired test, two-sided). The color of the arrow represents the average FC among genes experiencing a given transition. Transitions characterized by no significant changes in expression FC (Benjamini-Hochberg FDR ≥ 0.05) are represented by gray arrows. Upper panel: transitions from weaker to stronger active chromatin marking. Lower panel: transitions from stronger to weaker active chromatin marking. **d**: Examples showing different HMM states along transdifferentiation. For each gene, expression and chromatin tracks from one biological replicate are displayed, as well as normalized line plots averaging the signal from the two replicates. Profiles of HMM states for the three genes are shown at the bottom. Left panels: example of an up-regulated gene (*NUCB1*) with a constant HMM state profile along transdifferentiation. Middle panels: example of an up-regulated gene (*CCR5*) transitioning first from absence of marking state (a) to low marking state (b), and from this to active marking state (d). Right panels: example of a down-regulated gene (*MCAM*) transitioning from active marking state (d) to bivalent marking state (c).

Of the remaining genes, the vast majority (90%) go over just one-state transition during transdifferentiation. When considering DE genes, these transitions are generally associated with the expected transcriptional changes (Fig. 2c). Transitions from weaker to stronger active chromatin marking are accompanied by increases in gene expression (Fig. 2c, upper side; Fig. 2d, middle panels), while transitions from stronger to weaker active chromatin states are accompanied by decreases in gene expression (Fig. 2c, lower side; Fig. 2d, right panels). However, while transitions from active to strong active marking states (and vice versa) are more numerous, the corresponding fold changes in gene expression are lower, compared to transitions from low marking to active marking states (and vice versa). We observed activating transitions from the absent state mainly to the low marking state, further supporting the fact that marking by H3K4me1 and H3K4me2 is a prerequisite for the deposition of any other active histone modifications. On the other hand, we did not observe transitions from the strong active marking state to absence of marking, suggesting that the erasing of chromatin marks is not as an efficient process as its deposition.

Analysis of individual histone marks confirmed the HMM results. We determined whether the marks’ signals are stable or variable over time, analogously to what was done for gene expression profiles. The majority of genes present, indeed, stable chromatin profiles during transdifferentiation, even when focusing only on the differentially expressed ones (Table S3, left side; Fig. 3a). Lysine acetylation (H3K27ac and H3K9ac) is the most dynamic signal (Table S3, left side). Still, around 35% of DE genes show no changes in histone acetylation, despite being marked. Unexpectedly, only 8.5% of DE genes show changes in H3K27me3 throughout the process, although roughly half of them are down-regulated. Conversely, for a smaller number of silent and stably expressed genes we observed significant variations in their chromatin profiles over time (Table S4, Figs. 3b-c), comparable or even larger than for DE genes (Fig. S3c), although no changes could be detected in their expression profiles. We observed, in general, that differentially marked genes display clearer transdifferentiation trajectories compared to genes that are stably marked (Fig. S3d), further supporting that the contribution of gene expression and chromatin marks to cell state is not fully overlapping.

**Figure 3:**
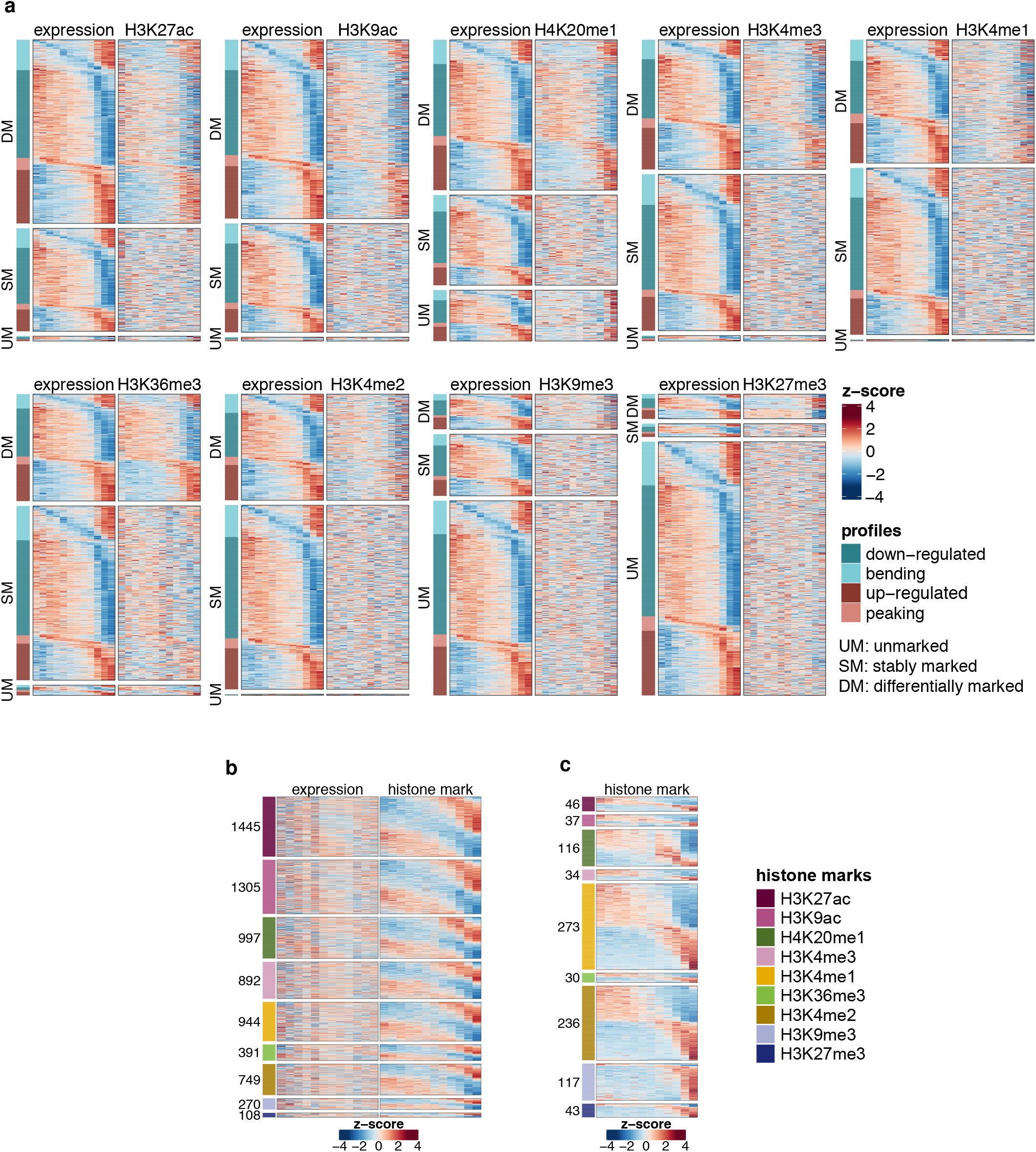
Uncoupling of expression and chromatin marks throughout transdifferentiation —. See also Fig. S3, Tables S3-4. **a**: Expression and chromatin profiles across the 12 time-points (columns) for the set of 8,030 DE genes, distinguishing between differentially marked (DM), stably marked (SM) and unmarked (UM) genes (rows). The profiles consist of row-normalized z-scores, computed independently for expression and chromatin marks. **b**: Expression and chromatin profiles over the 12 time-points (columns) for the set of stably expressed genes that are differentially marked for a given histone modification along transdifferentiation. The profiles consist of row-normalized z-scores, computed independently for expression and chromatin marks. The largest numbers of significantly variable profiles are observed for H3K27ac and H3K9ac. **c**: analogous representation to Fig. 3b for silent genes. In this case, H3K4me1 and H3K4me2 are the most variable marks throughout the process.

### Chromatin marking is associated with expression specifically at the time of gene activation

The limited number of chromatin HMM states indicates a coordinated behaviour of histone modifications. To investigate this behaviour at the resolution of individual marks and how it relates to gene expression, we first determined the type of association between each mark and expression along transdifferentiation, for each of the 8,030 genes that are differentially expressed (labels: unmarked, stably marked, positively correlated, uncorrelated and negatively correlated; see Fig. 4a, Table S3 and Methods). Then, we clustered the combinations of marks and types of association, and found that, in general, in a given gene, most marks show indeed the same type of association with expression (Fig. 4b). When clustering the genes based on these combinations, we found essentially three major groups (Figs. 4c, S4a). The first and largest cluster includes 4,995 DE genes (62%), presenting either stable or uncorrelated profiles for the majority of active marks, and absence of marking for H3K27me3 and H3K9me3 (Figs. 5a-b, upper panels). The second cluster includes 2,993 DE genes (37%), showing the canonical positive correlation between expression and most active modifications. A large proportion of these genes lack repressive marks, but a few of them (9%) exhibit the expected negative correlation with H3K27me3 (Figs. 5a-b, middle panels). Finally, the third and smallest cluster includes 102 genes (1%) characterized by an overall absence of both active and repressive marking, with the exception of H3K4me1 and H3K4me2 (Figs. 5a-b, lower panels).

**Figure 4:**
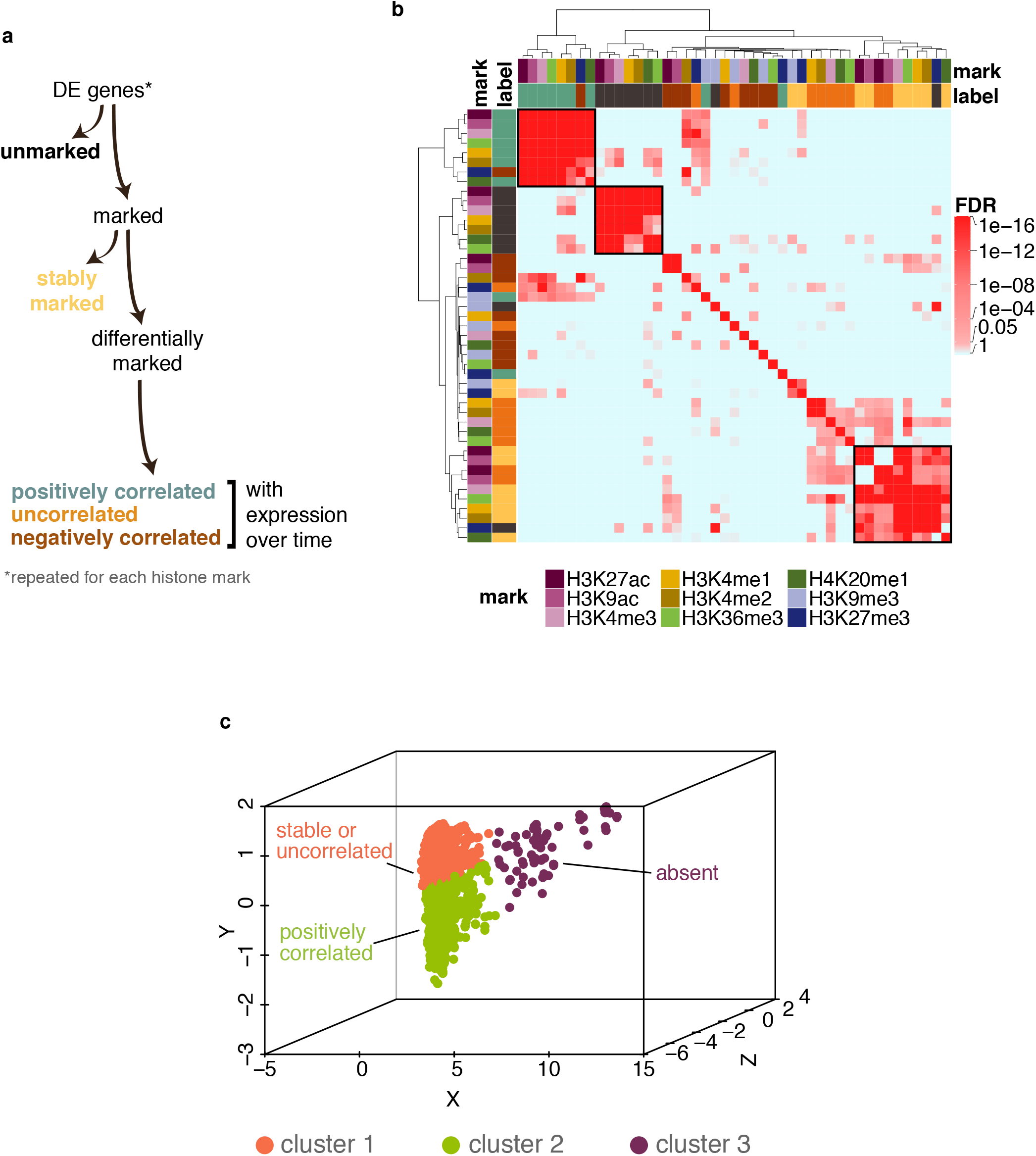
Chromatin marks show a coordinated behavior along transdifferentiation —. See also Fig. S4, Table S3. **a**: Decision-tree approach to label each of the 8,030 DE genes based on their chromatin marking status and its relationship with the expression profile over time. The approach is applied independently for each of the nine histone marks. The first branch distinguishes between unmarked (absence of peaks across all twelve time-points) and marked (presence of peaks in at least one time-point) genes. Within the set of marked genes, it further distinguishes between stably and differentially marked genes, i.e. genes characterized by absence and presence, respectively, of significant (maSigPro Benjamini-Hochberg FDR < 0.05) changes in chromatin signal along the process. Differentially marked genes are further classified into genes with positive, null or negative time-course correlation with expression. **b**: We assessed the overlap between sets of genes corresponding to the decision-tree labels across different histone marks (hypergeometric test). Hierarchical clustering of the FDR values identifies three main clusters: a) genes showing expression profiles positively correlated with H3K27ac, H3K9ac, H3K4me3, H3K36me3, H3K4me1, H3K4me2, H4K20me1, and negatively correlated with H3K27me3; b) genes unmarked for H3K27ac, H3K9ac, H3K4me3, H3K4me1, H3K4me2, H4K20me1 and H3K36me3; c) genes with stable or uncorrelated profiles for H3K27ac and H3K9ac, stable profiles for H3K4me3, H3K36me3, H3K4me1, H3K4me2, H4K20me1, and unmarked for H3K27me3. The color code for the labels is analogous to Fig. 4a. **c**: Similar results are obtained with Cluster Correspondence Analysis, a method that combines dimension reduction and cluster analysis for categorical data. Three-dimensional representation of the genes (analysis objects), grouped into three clusters (color-coded) based on the combinations of histone marks and labels they display.

**Figure 5:**
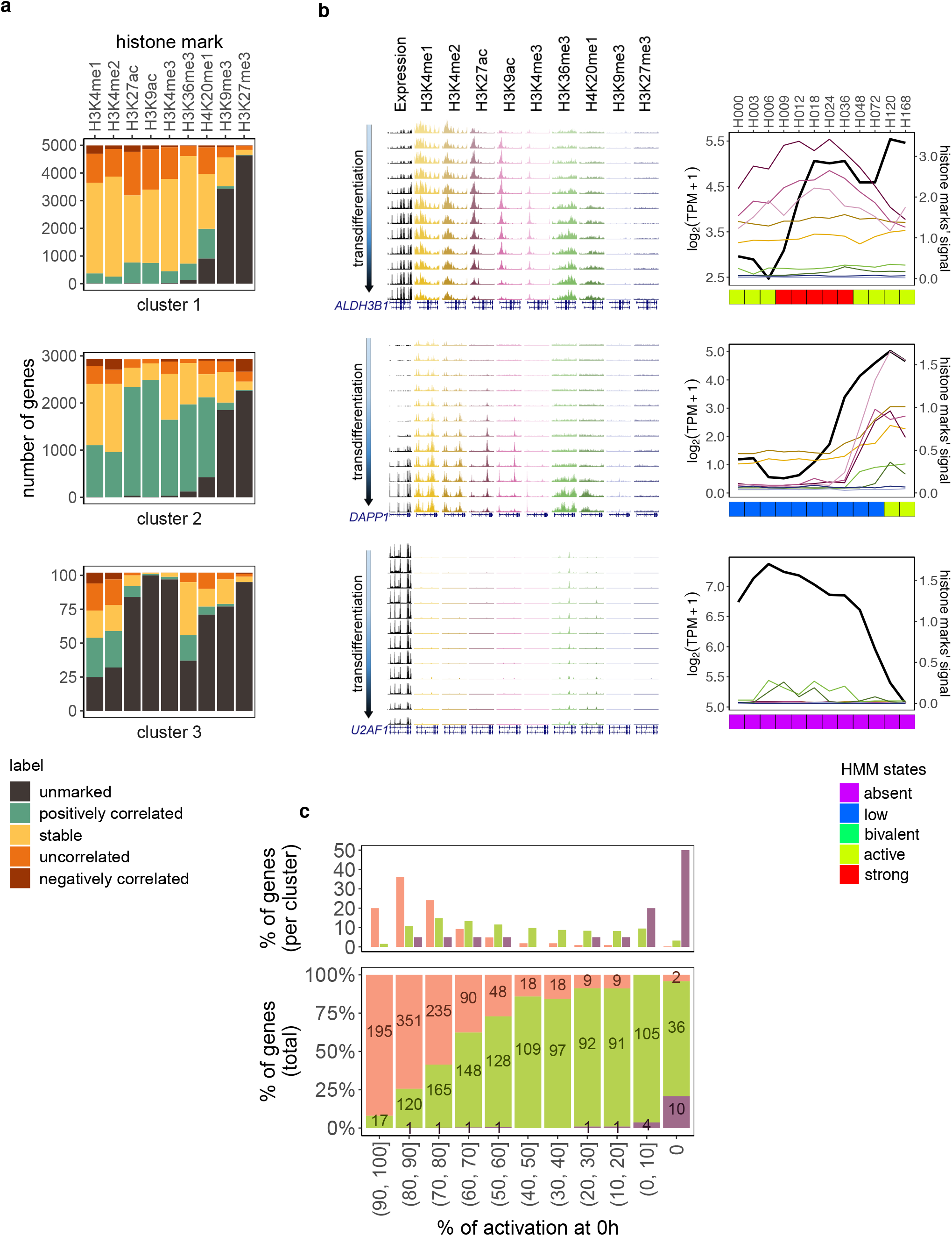
Chromatin marking is associated with expression specifically at the time of gene activation —. See also Fig. S4, Tables S5-6. **a**: Percent stacked bar plot representing, for each of the three clusters, the proportion of unmarked, stably marked, positively correlated, uncorrelated, and negatively correlated genes identified with respect to each histone mark. **b**: Examples of genes belonging to each cluster. For each gene, expression and chromatin tracks from one biological replicate are displayed, as well as normalized line plots averaging the signal from the two replicates. Profiles of HMM states for the three genes are shown at the bottom. Upper panels: example of an up-regulated gene (*ALDH3B1*) showing stable and uncorrelated profiles for active marking and absence of H3K9me3 and H3K27me3 along transdifferentiation. Middle panels: example of an up-regulated gene (*DAPP1*) showing positively correlated profiles for active marking and absence of H3K9me3 and H3K27me3 along transdifferentiation. Lower panels: example of a down-regulated gene (*U2AF1*) showing absence of marking along transdifferentiation. **c**: Percent stacked bar plot reporting the proportion of up-regulated genes in clusters 1-3 characterized by decreasing degrees of gene expression activation (bins of 10% decrement) at time-point 0h p.i. The degree of gene expression activation is defined as the ratio between the gene’s expression level at 0h and its maximum expression level along transdifferentiation.

Especially in the case of up-regulated genes, these clusters mostly reflect the level of gene activation when transdifferentiation starts (Figs. 5c, S4b-c). Genes in cluster 1 are already activated at the beginning of transdifferentiation, genes in cluster 2 are in early stages of activation or are activated early during transdifferentiation, while genes in cluster 3 are activated late during the process. The functions of the genes in these clusters are consistent with their level of activation at the beginning of transdifferentiation (Figs. S4d-e). In particular, genes in cluster 3 are associated with macrophage-specific functions, and we have found them lowly expressed and lowly marked in other cell types but CD14+ monocytes (Figs. S4f-g). Down-regulation of gene expression, on the other hand, appears to be largely uncoupled from chromatin changes, since most genes decreasing expression belong to cluster 1 (Fig. S4h).

### Gene expression changes anticipate changes in most active marks for up-regulated genes

The results above are suggestive that the association between gene expression and histone modifications occurs preferentially in a limited window of time during the initial stage of gene activation. Thus, to investigate the relationship between expression and chromatin marking precisely at this stage, we focused on the set of 257 up-regulated genes that are not expressed at 0 hours p.i., and that are, therefore, specifically activated during transdifferentiation. The vast majority of these genes (230, 89%) belong to cluster 2, that is, they are indeed characterized by positive correlation between gene expression and active chromatin marks. They are mostly associated with low and bivalent marking HMM states and, in 25% of the cases, transition into stronger marking states towards the end of transdifferentiation (Fig. S5a, upper panel).

To investigate the temporal relationship between gene activation and chromatin marking, for each up-regulated gene and histone mark we rescaled the expression and chromatin time-series profiles to the same range (0-100%), and identified the first time-point at which the expression level and the chromatin signal reach at least 25%, 50%, 75% and 100% (Fig. S5b). In this way, we determined whether active chromatin marking anticipates, co-occurs with, or follows gene expression. In contrast to the prevalent view, we did not find that most active marks anticipate activation of gene expression. At the first stage of up-regulation (25%), only marking by H3K4me1, H3K4me2 and H3K27ac anticipates more often than follows activation of gene expression (Figs. 6a-b), whereas for the other marks most changes follow expression up-regulation. These differences are progressively lost towards the end of the process (Figs. 6a, S5c).

**Figure 6:**
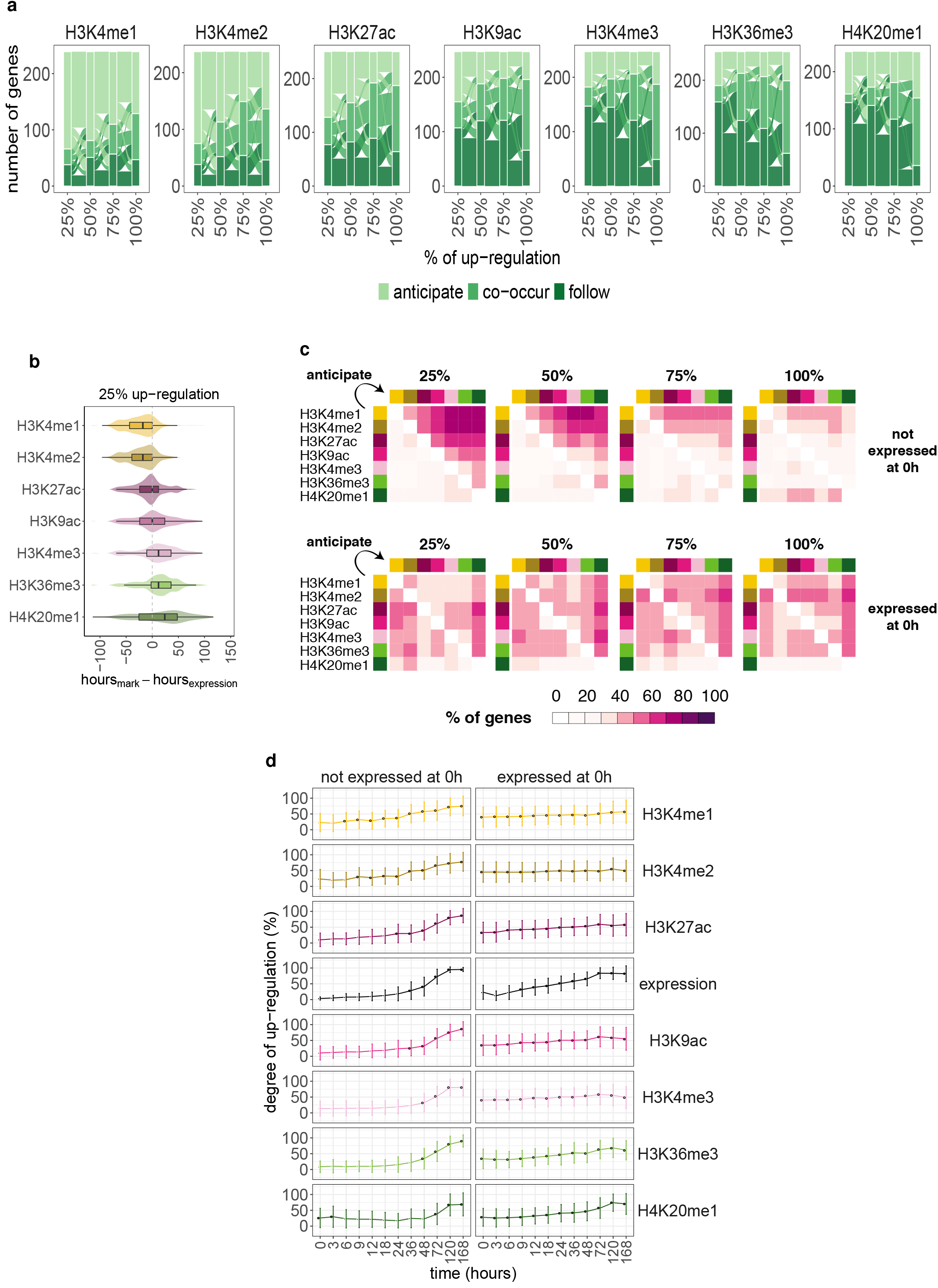
Gene expression changes anticipate changes in most active marks for up-regulated genes. —. See also Fig. S5. **a**: Alluvial plot describing, for each of the seven canonical active histone marks, the number of genes, out of 257 genes activated during transdifferentiation (i.e. up-regulated genes not expressed (< 1 TPM) at 0 hours p.i.), for which the up-regulation in a given mark’s signal anticipates (light green), co-occurs with (green) or follows (dark green) gene expression up-regulation. For more details see Fig. S5b. The flow lines indicate the number of genes exchanged among the three groups across increasing degrees of up-regulation. **b**: Lag (hours) between 25% up-regulation in histone marks’ signal and expression level for the 257 selected up-regulated genes. Negative lags correspond to changes in chromatin marks anticipating changes in gene expression; positive lags correspond to changes in chromatin marks following changes in gene expression. **c**: Upper panel: Heatmaps reporting the proportion (%) of genes activated during transdifferentiation whose changes in the chromatin mark on row *i* anticipate changes in the chromatin mark on column *j*. Like in the previous analyses, we considered four subsequent degrees of up-regulation (25%, 50%, 75% and 100%). e.g. the fraction reported in cell [row 1, column 2] of the first heatmap (25%), corresponds to the percentage of genes for which the 25% up-regulation in H3K4me1 signal (yellow - row 1) anticipates the 25% up-regulation in H3K4me2 signal (ochre - column 2). Lower panel: analogous to upper panel for the 629 up-regulated genes already expressed (> 25 TPM) at 0h p.i. For this latter set of genes there is not a precise order of increase in chromatin marks. **d**: Mean and standard deviation of time-series expression and chromatin profiles for the 257 (left panel) and 629 (right panel) up-regulated genes that are not expressed and highly expressed, respectively, at 0 hours p.i. The expression and histone marks’ time-series profiles of each gene were re-scaled to a 0-100% range prior to the analysis. We highlight in black the time-points at which the mean value is ≥ 25%.

To further decipher the precise order in which active chromatin signals are established over time, we computed, for a given mark, the fraction of genes whose changes either anticipate (Fig. 6c, upper panel) or co-occur with (Fig. S5d, upper panel) changes in each of the other six marks. When considering 25% of up-regulation, we observed that, in general, no marks anticipate H3K4me1, indicating that it is the first mark to increase, followed by H3K4me2 and H3K27ac (Fig. 6c, upper panel). This is consistent with the HMM analysis, which suggested that marking by H3K4me1 and H3K4me2 is a prerequisite for marking by other histone modifications (Fig. 2a). Changes in H3K4me1, H3K4me2 and H3K27ac most frequently precede increases in H3K9ac and H3K4me3. In all the comparisons, H3K36me3 and H4K20me1 follow the other marks (Fig. 6c, upper panel). As observed for gene expression, this precise order of marks’ deposition is progressively lost along transdifferentiation (Figs. 6c upper panel, S5d upper panel). Overall, this suggests that the deposition of active chromatin modifications follows a precise order at the time of initial gene activation (H3K4me1 > H3K4me2 > H3K27ac > expression > H3K9ac > H3K4me3 > H3K36me3 > H4K20me1; Fig. 6d left panel).

We performed a similar analysis with the set of 629 up-regulated genes that are already substantially expressed at 0 hours p.i. (> 25 TPM). These genes belong mostly to cluster 1 (389, 62%), that is, their expression profiles are uncoupled from changes in chromatin marking, and they actually remain in active chromatin states during transdifferentiation (Fig. S5a lower panel). For these genes we did not find preservation in the pattern of chromatin deposition with respect to expression (Fig. S5e), nor in the deposition of the marks (Figs. 6c lower panel; Fig. 6d right panel; S5d lower panel).

### A model to explain the coupling between transcription and chromatin marking over time

Altogether, our results show that the canonical association between histone modifications and gene expression mainly occurs in a limited window of time preceding and following initial gene activation. We specifically propose a model (Fig. 7a) in which the activation of gene expression is anticipated by deposition of H3K4me1, H3K4me2 and, less frequently, of H3K27ac at promoter regions. The deposition of other marks typically enriched either at promoters (H3K9ac, H3K4me3) or over the gene body (H3K36me3, H4K20me1) is concomitant to or, more often, follows (and may be induced by) gene activation. After this initial stage of gene activation, further changes in gene expression, comparable or even stronger, appear to be mostly uncoupled from changes in histone modifications (Fig. 7b, compare left and right panels).

**Figure 7:**
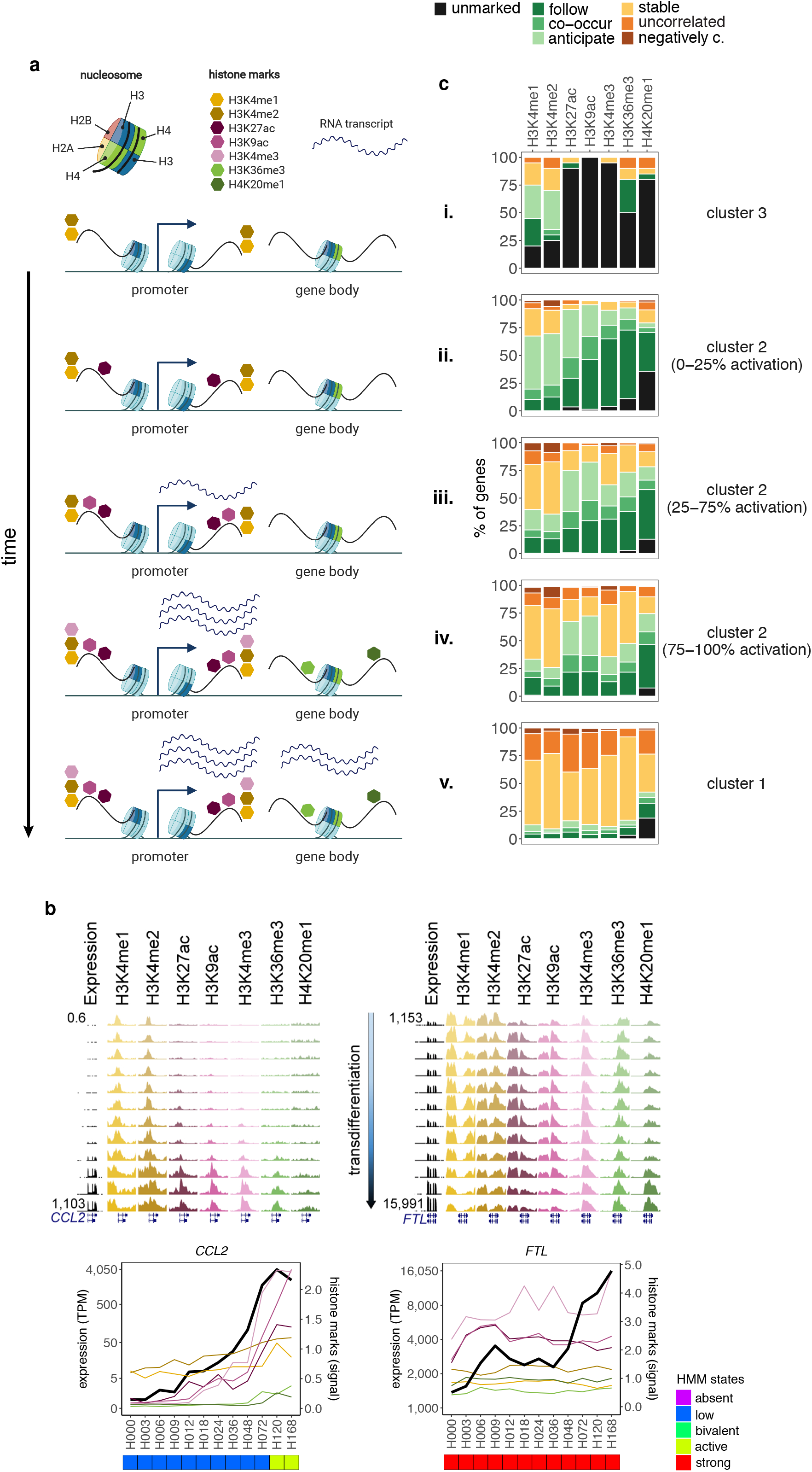
A model to explain the coupling between transcription and chromatin marking over time. **a**: According to our model, chromatin marking correlates with expression specifically during the first stage of gene activation, and the deposition of histone marks follows a specific order. Further changes in gene expression that happen later in time are mostly uncoupled from chromatin marking. **b**: Examples of up-regulated genes inactive (*CCL2*) and highly active (*FTL*) at the beginning of transdifferentiation. For each gene, expression and chromatin tracks from one biological replicate are displayed, as well as normalized line plots averaging the signal from the two replicates. Profiles of HMM states for the two genes are shown at the bottom. Left panels: for *CCL2*, most active histone modifications follow gene activation, with the exception of H3K4me1 and H3K4me2, which anticipate it. Right panels: for *FTL*, most active histone modifications remain stable along transdifferentiation, even though its absolute increase in expression is much higher than that of *CCL2*. **c**: Percentage (%) of unmarked, stably marked, positively correlated, uncorrelated and negatively correlated profiles within cluster 3, cluster 2 (0-25%, 25-75%, 75-100% activation level), and cluster 1 up-regulated genes. Positively correlated genes are further separated into genes whose histone mark’s up-regulation anticipates, co-occurs with or follows gene expression up-regulation.

This model explains our observations well. The patterns of association between chromatin marking and gene expression (as defined in Fig. 4a) for genes in different degrees of activation when transdifferentiation starts (0h p.i.) reflect how this association changes as gene activation proceeds (Fig. 7c). Up-regulated genes that are silent when transdifferentiation starts (mostly in cluster 3) lack almost all “activating” histone modifications, possibly with the exception of H3K4me1 and H3K4me2 (i.). Up-regulated genes in cluster 2 that are lowly or not activated at 0h show mostly correlated patterns of expression and chromatin marking. In these genes, most marks, with the exception of H3K4me1, H3K4me2 and H3K27ac, follow rather than anticipate expression (ii., see also Fig. 7b, left panel). As we consider genes with increasing degrees of activation at 0h (and thus, in increasingly advanced states of activation), the fraction of genes with correlated patterns of expression and chromatin marking decreases, while the fraction of genes with stable or uncorrelated chromatin profiles (iii. and iv.) proportionally increases. The temporal order of activation of marks observed in early activation stages is also gradually lost. Finally, for genes in cluster 1 (v.), which are already highly active when transdifferentiation starts, changes in gene expression, even if substantial, are mostly uncoupled from chromatin marking, showing uncorrelated or stable profiles (see also Fig. 7b, right panel).

## Discussion

Epigenetics was initially defined as “the branch of biology that studies the causal interactions between genes and their products which bring the phenotype into being” (Waddington, 1942). In a more contemporary definition, “an epigenetic trait is a stably heritable phenotype resulting from changes in a chromosome without alterations in the DNA sequence” (Berger et al., 2009). The epigenetic mechanisms leading to the development of an individual or to the differentiation of a cell lineage from the unique genotype of the organism have been largely studied during decades. Although initial references to the mechanisms by which epigenetics promotes cell memory and leads cell fate did not relate to its ability to regulate gene expression, a causative role for epigenetic modifications in controlling transcription has been later pointed out (see Bannister and Kouzarides, 2011, and Rivera and Ren, 2013, for reviews about different aspects related to epigenetics and its role in regulating gene expression), and it has even been shown that some epigenetic features, such as histone modifications, are accurate predictors of gene expression (Karlic et al., 2010; Dong et al., 2012; Sekhon et al., 2018) and the other way around (Yin et al., 2019).

However, the causal/consequential relationship between chromatin modifications and gene expression represents a long-standing discussion (Henikoff and Shilatifard, 2011), and a number of reports have challenged the causal role that has been broadly attributed to chromatin modifications (Dorighi et al., 2017; Rickels et al., 2017; Krogan et al., 2003; Hödl and Basler, 2012). Still, and despite the efforts dedicated to this problem and the vast literature produced, the actual relationship between histone modifications and the regulation of gene expression remains unsolved.

This is partially due to the few available studies in which gene expression and histone modifications have been both consistently monitored through time in a given dynamic system. Differentiation models are suitable to study the relationship between gene expression and chromatin marking, as they provide a dynamic system that allows to decipher the order of the events. In this work, we have used the transdifferentiation of BLaER1 cells (pre-B cells) into macrophages, a model that has proven to be highly efficient (Rapino et al., 2013), and we have generated high-quality data on the transcriptome and the epigenome in twelve time-points along the seven days the transdifferentiation process lasts. Our analysis of these data has uncovered some fundamental features of chromatin organization in human genes and of the relationship between gene expression and histone modifications.

Our analyses have also contributed to a better understanding of the molecular events underlying transdifferentiation of pre-B cells into macrophages. Despite the fact that, to our knowledge, there is no retro-differentiation during the process (Di Tullio et al., 2011; Rapino et al., 2013), the joint PCA of gene expression and chromatin marks suggests that BLaER1 cells undergo an intermediate state (Fig. 1c). This intermediate state is characterized by chromatin changes not accompanied by changes in gene expression (Fig. S6), and vice versa by changes in gene expression not associated with chromatin changes (Fig. S6a). Although it is often assumed that the transcriptome is the main determinant of cell state, these results suggest that epigenetic modifications contribute to cell state in a manner that cannot be fully recapituted by gene expression. Thus, neither the epigenome nor the transcriptome can be fully predictive of one another.

Consistently, we found that the association between gene expression and chromatin modifications is overall weaker than reflected by the correlations reported so far, which have been mostly computed in a particular steady-state cellular condition (Fig. 1d). These artifactually strong correlations result from the largely constrained nature of the human epigenome and transcriptome. In particular, a large fraction of genes in the human genome (likely more than 50%, Pervouchine et al., 2015) are either invariably silent and not marked, or expressed and marked across most cellular states. Genes with stable epigenomes and transcriptomes drive the correlations to large values when computed in a particular cell condition, and explain why models relating gene expression to histone modifications inferred in a particular cell type have high predictive power in other cell types (Karlic et al., 2010; Dong et al., 2012; Sekhon et al., 2018; Yin et al., 2019), even though there is no true causality involved in the relationship between chromatin and expression. The steady-state correlations represent an example of the Sympson’s paradox (Simpson, 1951), by which the data can show different or even opposite behavior if subgroups within the dataset are considered.

HMMs have been widely used to summarize patterns of combinations of multiple histone modifications into a limited number of chromatin states. However, in most cases so far, they have been used to segment the genome sequence (Ernst and Kellis, 2012; Hoffman et al., 2012; Song and Chen, 2015; Zhang et al., 2016; Zhang and Hardison, 2017). Here, instead, we used them, we believe for the first time, to segment time along a dynamic differentiation process. The HMM segmentation reveals that, even though the number of possible histone combinations is very large (if nine histones are considered, 2^9^ = 512 combinations are possible), most genes are actually found in one among only about five major states (Fig. 2a). This challenges to some extent the notion of a histone code (Strahl and Allis, 2000). Further supporting the limited number of genic chromatin states, we found that marks act in a coordinated manner, meaning that genes showing a stable profile for one histone modification tend also to present stable profiles of the other marks, and that genes showing absence of one active mark tend to be void of all positive modifications (Figs. 4b-c, S4a). Most genes remain in the same chromatin state during transdifferentiation, irrespective of whether they are or not differentially expressed, explaining the low correlation between gene expression and chromatin marks throughout time. Analysis of individual histone modifications further uncovered two unexpected findings regarding the chromatin marks typically associated with gene silencing. First, we observed that, although roughly 4,000 genes are down-regulated during the process, only 10% of them present H3K27me3 marking in at least one time-point, indicating that the majority of genes that are silenced along transdifferentiation do not depend on Polycomb repression. Second, we saw that H3K9me3 marking at transcription start sites is associated more frequently with active transcription than gene silencing (see Table S3, right side), contrary to what has been previously reported (Ninova et al., 2019). Actually, H3K9me3 at the transcription start site has been previously related to active expression in malignant cells (Wiencke et al., 2008). Furthermore, these analyses also allowed us to identify a number of silent or stably expressed genes along transdifferentiation that show changes in chromatin marking (Figs. 3b-c).

While there is a general lack of coupling between gene expression and chromatin marking, there is a temporal relationship between gene expression and the different histone modifications at the time of gene activation. We propose a model (Fig. 7a) in which activation of gene expression is anticipated by deposition of H3K4me1, H3K4me2, while deposition of other marks is concomitant or, more often, follows gene activation, being the gene body marks the last ones to be incorporated. The order of chromatin marking in our model is in agreement with the observed deposition of histone modifications upon induction of gene expression in human melanoma cells (Rybtsova et al., 2007), and with the notion that the methylation of some histone residues depends on the transcription machinery (Krogan et al., 2003). While we observed that certain modifications, such as H3K4me1/2 and H3K27ac tend to anticipate gene expression, this does not necessarily mean that they are the cause of transcription initiation. Actually, we have also observed particular cases in which these marks are deposited post-activation (for an example see Fig. 5b, middle panels). After the initial stage of gene activation, further changes in gene expression, even if substantial, appear to be mostly uncoupled from changes in histone modifications (Fig. 7b). It is tempting to speculate that after the initial burst of transcription, histone residues are saturated with modifications, and that therefore, any further up-regulation of gene expression cannot possibly be accompanied by increased levels of histone modifications.

We do have identified a small set of genes that are expressed in the absence of any histone modification, with the exception of H3K4me1 and H3K4me2 (Figs. 4c, 5a-b lower panels). A few of these are activated later during the transdifferentiation process, and therefore we lack the temporal resolution to detect post-activation marking. Still, many of these genes are down-regulated or stably expressed, and are unmarked even at the beginning of transdifferentiation (for an example see Fig. 5b, lower panels). Gene activation without histone modifications has been previously observed for developmentally regulated genes in the fruit fly (Pérez-Lluch et al., 2015).

Here we have focused specifically on the dynamics of chromatin modifications during up-regulation. Our results suggest that down-regulation appears to be largely uncoupled from chromatin changes (Fig. S4h). However, while RNA sequencing-inferred expression levels can be used to approximately identify the time at which a gene is initially activated, differences in RNA stability may confound the identification of the time-point at which a gene is fully inactivated. Indeed, RNAs can be detected long after gene inactivation, for a time likely to be specific to each individual gene. Therefore, the data that we have generated does not have the appropriate resolution to discard that this lack of coupling during down-regulation is partially caused by the difficulty in precisely identifying the time-point at which genes stop being expressed.

The multi-omics data that we have generated during the pre-B cell transdifferentiation into macrophages has allowed us to address with unprecedented resolution some fundamental questions regarding the dynamics of chromatin marking and gene expression during cellular differentiation, and have contributed to shed light on some long-standing questions in the field. Further mining of this data resource will certainly contribute to a deeper understanding of the epigenetic layer of gene regulation.

## Acknowledgments

We thank Thomas Graf and Francesca Rapino for donating BLaER1 cells and for helpful discussions. We thank Sebastian Ullrich, Carme Arnan and Vasilis Ntasis for helpful discussion about the data. We thank Montserrat Corominas, Guillaume Filion and Luciano Di Croce for insightful suggestions. We thank Diego Garrido-Martín, Manuel Muñoz and Javier Martín-Vallejo for statistical advice. We also thank the Genomics, the Flow Cytometry and the Bioinformatics Core Units of the CRG (Barcelona, Spain). We thank Romina Garrido for administrative support. We thank the ENCODE Consortium, in particular Thomas Gingeras’, Bradley Bernstein’s, John Stamatoyannopoulos’ and Peggy Farhnam’s laboratories, for data production. This work was performed under the financial support of the European Community under the FP7 program (ERC-2011-AdG-294653-RNA-MAPS). B.B. is supported by the fellowship 2017FI_B00722 from the Secretaria d’Universitats i Recerca del Departament d’Empresa i Coneixement (Generalitat de Catalunya) and the European Social Fund (ESF). C.C.K. is supported by the CERCA Programme / Generalitat de Catalunya and FEDER under project VEIS-001-P-001647. B.R.C. is supported by the Ministerio de Ciencia, Innovación y Universidades de España under grant FJCI-2017-34353. We also acknowledge Agencia Estatal de Investigación (AEI) and FEDER under project PGC2018-094017-B-I00. All authors acknowledge the support of the Ministerio de Ciencia, Innovación y Universidades de España to the EMBL partnership, the Centro de Excelencia Severo Ochoa, and the CERCA Programme / Generalitat de Catalunya.

## Author Contributions

R.G. and R.J. conceived the project. B.B., S.P-L. and R.G. designed the study. B.B. performed the computational analyses. A.A. performed the ChIP-seq experiments. A.E. and M.S. performed the RNA-seq experiments. C.C.K. and E.P. contributed to data quality check and processing. C.C.K., R.N., M.R-R. and B.R.C. contributed tools and ideas to perform experiments and computational analyses. B.B., S.P-L. and R.G. wrote the manuscript with the contribution of all authors.

## Declaration of Interests

The authors declare no competing interest.

## STAR Methods

### RESOURCE AVAILABILITY

#### Lead Contact

Further information and requests for resources and reagents should be directed to and will be fulfilled by the Lead Contact, Roderic Guigó.

#### Materials Availability

This study did not generate new unique reagents.

#### Data and Code Availability

The code generated during this study is available at https://github.com/bborsari/Borsari_et_al_transdifferentiation_chromatin. A complete list of scripts used for each analysis described in the section *Method details* can be found at https://github.com/bborsari/Borsari_et_al_transdifferentiation_chromatin/blob/master/bin/table.scripts.tsv. When not specified in the text, the code used for a given analysis is included in the corresponding figure’s script.

RNA-seq and ChIP-seq raw and processed data from this study have been submitted to ArrayExpress (https://www.ebi.ac.uk/arrayexpress/) under accession numbers E-MTAB-9790 and E-MTAB-9825, respectively.

Processed data in GRCh38/hg38 assembly from this study is available for visualization at the UCSC Genome Browser (Tyner et al., 2017, http://genome.ucsc.edu/). The track data hub is available at https://public-docs.crg.es/rguigo/Data/bborsari/hubs/ERC_human_hub/hub.txt.

A web page has also been implemented to gather all information regarding the Chromatin and Transcriptomics Dynamics Project (http://rnamaps.crg.eu/). The web page provides information about all experiments and replicates performed during the project, as well as access to the data in ArrayExpress and the UCSC Genome Browser.

ENCODE data is freely available on the ENCODE portal (https://www.encodeproject.org/). Experiments and files accession IDs for RNA-seq and ChIP-seq data are reported in Supplementary Tables S5 and S6, respectively.

### EXPERIMENTAL MODEL AND SUBJECT DETAILS

#### Transdifferentiation of BLAER1 cells to macrophages

For the transdifferentiation process we made use of the Burkitt lymphoma cell line BlaER1, as described in Rapino et al., 2013. Induction of transdifferentiation (treatment with 100 μM **β**-estradiol and growth in the presence of 10 nM Il-3 and 10 nM CSF-1) has been described in Bussmann et al., 2009, and Xie et al., 2004. The process was monitored at 12 time-points (as described in Rapino et al., 2013): 0, 3, 6, 9, 12, 18, 24, 36, 48, 72, 120 and 168 hours post-induction (p.i.; Fig. 1a).

### METHOD DETAILS

#### RNA-seq library preparation and sequencing

Two independent biological replicates for each time-point were performed. Briefly, cells were lysed with QiAzol (Qiagen, The Netherlands). Chloroform was added to each sample, and RNA contained in the aqueous solution was isolated and purified by using RNeasy mini kit columns (Qiagen, The Netherlands). Poly A+ libraries were prepared with 1 μg of total RNA and using TruSeq Stranded mRNA Library Prep Kit (Illumina, USA) according to the manufacturer’s protocol. Libraries were analyzed using Agilent DNA 1000 chips to determine the quantity and size distribution, and sequenced paired-end 75-bp on an Illumina HiSeq 2000.

#### ChIP-seq library preparation and sequencing

ChIP-seq experiments of nine histone marks (H3K4me1: Abcam ab8895; H3K4me2 : Millipore 07-030; H3K4me3: Abcam ab8580; H3K9ac: Abcam ab4441; H3K27ac: Diagenode C15410192; H3K36me3: Abcam ab9050; H4K20me1: Abcam ab9051; H3K9me3: Abcam ab8898; H3K27me3: Millipore 07-449) were performed in two independent biological replicates for each time-point. Cells were crosslinked with formaldehyde 1% (Sigma) for 10’ at room temperature. The reaction was stopped by adding glycine to 0.25 M final concentration for 10’ at room temperature. Fixed cells were resuspended in 100 μL of lysis buffer (SDS 1%, EDTA 10 mM, TrisCl 50 mM and protease inhibitors). The lysate was sonicated for 25’ using Covaris S2 system in TC12 tubes (Duty cycle 20%, Intensity 8, cycles/burst 200, water level 15). The cleared supernatant was used immediately in ChIP experiments or stored at −80 °**C**. 5 μg of sonicated chromatin were diluted in 900 μL RIPA buffer — H3K4me3, H3K9ac, H4K20me1, H3K27me3 and H3K27ac (140 mM NaCl, 10 mM Tris-HCl pH 8.0, 1 mM EDTA, 1% Triton X-100, 0.1% SDS, 0.1% Na deoxycholate, protease inhibitors) —, RIPA 2X — H3K4me1, H3K4me2 and H3K9me3 (280 mM NaCl, 10 mM Tris-HCl pH 8.0, 1 mM EDTA, 2% Triton X-100, 0.2% SDS, 0.2% Na deoxycholate, protease inhibitors) —, or RIPA 1X 1% triton — H3K36me3 (280 mM NaCl, 10 mM Tris-HCl pH 8.0, 1 mM EDTA, 1% Triton X-100, 0.2% SDS, 0.2% Na deoxycholate, protease inhibitors). For H3K4me3, H3K36me3, H3K9ac and H3K27me3 ChIPs, chromatin and antibodies were incubated overnight, rotating at 4 °**C** with 0.125-5 μg of specific antibody and samples were then incubated for 2 hours rotating at 4° **C** with Dynabeads protein A for immunoprecipitation (Invitrogen) to recover the bound material. For H3K4me1, H3K4me2, H3K9me3, H4K20me1 and H3K27ac ChIPs, antibodies were coated to protein A magnetic beads for 2 hours at 4 °**C** prior to overnight incubation with chromatin. In all cases, beads were washed for 10’ three times in 1 mL of the corresponding immunoprecipitation buffer without protease inhibitors, then washed once in 1 mL LiCl buffer (0.25 M LiCl, 0.5% NP-40, 0.5% sodium deoxycholate, 1 mM Na-EDTA, 10 mM Tris-HCl, pH 8.0), and finally washed twice in 1 mL of TE buffer (1 mM Na-EDTA, 10 mM Tris-HCl, pH 8.0). ChIPped material was incubated with DNase-free RNase at 50 μg/mL for 30’ at 37 °**C**. Chromatin was reverse-crosslinked by adding SDS (0.5% final concentration) and Proteinase K (500 μg/mL final concentration) and incubated overnight at 65 °**C**. ChIPped chromatin was then purified with Qiaquick PCR purification columns (Qiagen) following the manufacturer’s instructions. ChIP libraries were prepared with 1-5 ng of DNA and using NebNext Ultra DNA library prep kit for Illumina (New England Biolabs) according to the manufacturer’s protocol. Libraries were analyzed using Agilent DNA High Sensitivity chips to determine the quantity and size distribution, and sequenced single-read 50-bp on an Illumina HiSeq 2000.

In total, 264 samples were sequenced (24 by RNA-seq, 216 by ChIP-seq, 24 by ChIP input).

#### RNA-seq data processing and analysis

Data was processed using the *grape-nf* (https://github.com/guigolab/grape-nf) Nextflow (DI Tommaso et al., 2017) pipeline. RNA-seq reads were aligned to the human genome (assembly GRCh38, Gencode annotation version 24) using the STAR software version 2.4.0j (Dobin et al., 2013). We allowed a maximum number of mismatches equal to 4% of the read length. Only alignments for reads mapping to ten or fewer loci were reported. Quantification of genes and transcripts was done with RSEM version 1.2.21 (Li and Dewey, 2011). TPM calculation was performed after removing mitochondrial genes.

From the set of 19,831 protein-coding genes (Gencode v24), we selected 10,696 expressed genes with a maximum expression during transdifferentiation ≥ 5 TPM in both replicates, and 1,552 silent genes (0 TPM in all time-points and replicates). Based on this set of 12,248 genes, we quantile-normalized the expression matrices (log_2_-transformed TPM, pseudocount of 1) across replicates and time-points using the R package preprocessCore (Bolstad et al., 2003, script: `quantile.normalization.R`), and obtained the mean expression levels between replicates (script: `matrix_matrix_mean.R`).

To detect significant gene expression changes along transdifferentiation, we used the R package maSigPro (Nueda et al., 2014) with replicates handled internally. Function `p.vector()` was run with default parameters: `Q = 0.05, MT.adjust = “BH”, min.obs = 20` (script: `maSigPro.wrapper.R`). We defined as stably expressed those genes reporting a maSigPro FDR value ≥ 0.05 (*n* = 2,666).

As concerns the identification of up-regulated, down-regulated, peaking and bending genes, we performed a two-step classification across the 8,030 genes with significantly variable gene expression profiles. Briefly, we first focused on profiles with at least two-fold change (in *log_2_* scale this change corresponds to 1) and identified monotonic up-regulations and down-regulations; peaking profiles were defined as monotonic increases followed by monotonic decreases, bending profiles as the opposite (script: `classification.log2.pl`). All other significantly variable genes with fold-change < 2 were assigned to one of these four groups following hierarchical clustering (distance measure: *euclidean*; clustering method: *complete*; script: `classification.2.R`).

#### ChIP-seq data processing and analysis

Data was processed using the *ChIP-nf* (https://github.com/guigolab/chip-nf) Nextflow pipeline. ChIP-seq reads were aligned to the human genome assembly (GRCh38) using the GEM mapping software (Marco-Sola et al., 2012), allowing up to two mismatches. Only alignments for reads mapping to ten or fewer loci were reported. Duplicated reads were removed using Picard (http://broadinstitute.github.io/picard/). Pile-up signal from bigWig files was obtained running MACS2 (Zhang et al., 2008) on individual replicates. No shifting model was built. Instead, fragment length was set to 250 bp and was used to extend each read towards the 3’ end (using the `--extsize` option). Pile-up signal was normalized by scaling larger samples to smaller samples (using the default for the `--scale-to` option) and adjusting signal per million reads (enabling the `--SPMR` option). Peak calling was performed using Zerone (Cusco and Filion, 2016) with replicates handled internally, and passed the filter for all pairs of replicates (`advice: accept discretization`).

To check library complexity, we computed the fraction of non-redundant mapped reads (Landt et al., 2012) (recommended threshold: NRF ≥ 0.8) for each ChIP-seq experiment, and found a minimum NRF value of 0.92. Additionally, to evaluate the global ChIP enrichment, we computed the fraction of reads in peaks (Landt et al., 2012) (recommended threshold: FRiP ≥ 0.01), and found a minimum FRiP value of 0.05.

The intersection / overlap analyses described below were performed with the function `intersectBed` of BEDTools software v2.27.1 (Quinlan and Hall, 2010).

To select the genomic location enriched, on average, in a specific histone mark (region of interest), we focused on an up-stream and down-stream 5 Kb region (±5 Kb) with respect to the first annotated Transcription Start Site (TSS) of the gene, and retrieved 6,063 protein-coding genes that did not overlap any other gene body ±5 Kb. For each histone modification we then selected, among the 6,063 genes, those with peaks in the ±5 Kb promoter region in all the 12 time-points, and computed, using the function `aggregate` from the bwtool software (Pohl and Beato, 2014, script: `bwtool.aggregate.ChIPseq.sh`), the mean pile-up signal for each experiment. Based on this analysis, we decided to select as regions of interest i) the gene body for H3K36me3 and H4K20me1, ii) ±2 Kb with respect to the TSS for all other marks (Fig. S1c). A comprehensive catalogue of all non-redundant (same ensembl gene ID and start coordinate) TSSs annotated for the selected 12,248 in Gencode v24 was obtained with the script `non.redundant.TSS.sh`.

To compare expression and chromatin profiles over time, we quantified, for each of the nine histone marks, the amount of pile-up signal associated with a gene at each time-point (script: `get.matrix.chipseq.sh`). Briefly, if a peak was present in the region of interest of a gene at a specific time-point, we considered the mean pile-up signal in the intersection between the peak and the region of interest, otherwise we computed the mean pile-up value in the entire region of interest. In the presence of multiple peaks and/or multiple regions of interest (e.g. in case of multiple TSSs annotated for the same gene), we considered the highest of all observed values. Matrices of histone marks’ signals for the selected 12,248 protein-coding genes were quantile-normalized across replicates and time-points using the R package preprocessCore as done for gene expression. For all down-stream analyses, we used the mean signal between replicates.

#### Principal Component Analysis of expression and chromatin data

For this type of analysis we made use of the transposed expression and chromatin matrices generated as described in sections *RNA-seq data processing and analysis* and *ChIP-seq data processing and analysis*, respectively. Therefore, genes (columns) and time-points (rows) were used as variables and observations, respectively. We centered and scaled each of the ten transposed matrices independently, obtaining z-score profiles for each time-point monitored at expression and histone marks’ level. For the joint Principal Component Analysis (PCA) reported in Fig. 1c across expression and the nine histone marks, we included as variables the subset of 10,658 genes with non-missing (NA) *z*-score profiles in all ten matrices. As a consequence, 1,590 genes were excluded from this analysis, 98% of them being the silent genes (1,552). For the PCAs reported in Fig. S3d, we considered for each histone modification the corresponding sets of DE genes that are either stably or differentially marked.

#### Analysis of the degree of correlation between expression levels and chromatin signals

Steady-state correlations between gene expression levels and each histone mark’s signals were computed at individual time-points considering the entire set of 12,248 selected protein-coding genes. In this case, Pearson *r* measured the degree of correlation between the vector of 12,248 expression levels and the vector of 12,248 mark signals at a given time-point (Fig. 1d, dots). Time-course correlations were measured, instead, at the level of individual expressed genes. Silent genes were not considered for this analysis, because of the zero standard deviation in their time-series expression profile (i.e. 0 TPM in all time-points). Thus, for each gene and histone mark we obtained the Pearson *r* correlation coefficient between the vector of 12 expression levels (i.e. the expression levels measured at the 12 time-points) and the vector of 12 mark signals. The distributions of Pearson *r* correlation coefficients for the set of (differentially + stably) expressed genes are depicted with box plots and violin plots in Fig. 1d. Randomized steady-state and time-course correlation coefficients were computed as described above following a 1,000-permutations scheme on each histone mark’s matrix. Briefly, while we kept the original expression matrix, the columns (time-points) of the matrix corresponding to a given mark’s signal were permuted without repetition 1,000 times (for an example, see Fig. S2a, lower panel). In the case of steady-state correlations we report, for each expression time-point, the Pearson *r* averaged over 1,000 rounds of permutation of chromatin time-points (Fig. S2b, dots). In the case of correlations computed across time-points (time-course), we computed, for each gene, the Pearson *r* averaged over the 1,000 rounds of permutations. The distributions of the resulting coefficients across the set of expressed genes are depicted in Fig. S2b (box plots and violin plots). Correlations were computed with the R function `cor()`. Permutations without replacement of the chromatin time-points were performed consistently across histone marks with the R function `sample()`, by setting an independent seed for each round of permutations. The correlation values reported in Fig. S2c are an analogous exercise to Fig. 1d on the set of 8,030 differentially expressed genes.

#### Multivariate Hidden Markov Model analysis

A multivariate Hidden Markov Model (HMM) was fitted to the entire ChIP-seq dataset to approximate the set of underlying chromatin states reported by the 12,248 selected protein-coding genes along the transdifferentiation process. Specifically, we provided as input a matrix of dimensions 146,976 rows × 9 columns, which collected for each gene and time-point (12,248 genes, 12 time-points) the signal of each of the 9 histone marks after quantile normalization (for a description of these calculations see previous section *ChIP-seq data processing and analysis*). The collective behavior of the nine histone marks along the twelve time-points was modelled as an independent time-series for each gene, using Gaussian distributions. The model then reprocessed each gene’s data to estimate the chromatin state of each gene at each time-point, and provide a time series of chromatin states for each gene. HMM was performed using the R package depmixS4 (Visser and Speekenbrink, 2010), in particular functions `depmix(), fit()` and `posterior()` (script: `HMM.wrapper.marks.R`). We repeated the analysis for increasing numbers of states (between 2 and 20), and recorded the log likelihood of each model (the 20-states model reached the maximum number of iterations in EM without convergence). We found that somewhere between five and eight states approximate the elbow point of the log likelihood curve (Fig. S3a), and observed that the combinations of histone marks represented by five states were consistent with manual inspection of pile-up histone marks profiles in the UCSC genome browser. We thus set for five states. The response parameters of the nine histone marks corresponding to each of these states are reported in Fig. 2a. In this case, the *Intercept* values of each histone mark across the five states were re-scaled to a range 0-1 to enable the comparison among different states and marks. HMM sequence hierarchical clustering across the 12,248 genes was performed with the TraMineR (Gabadinho et al., 2011) and pheatmap (https://github.com/raivokolde/pheatmap) R packages (clustering distance: *euclidean*, clustering method: *Ward.D2*). The arc diagram representation in Fig. 2c was obtained with the R package arcdiagram (https://github.com/gastonstat/arcdiagram).

#### Decision-tree labelling

In the Methods section *ChIP-seq data processing and analysis* we introduced the distinction between genes with and without peaks of a given mark at a given point in the region of interest (gene body for H3K36me3 and H4K20me1; TSS ±2 Kb for all other marks). Following this first assessment, we classified as unmarked those genes that were consistently unmarked throughout the whole process of transdifferentiation, i.e. with no peaks called at any time-point in the region of interest. Conversely, marked genes reported peak calls of a given mark in the region of interest in at least one time-point (Fig. 4a).

Within the set of marked genes, we defined as stably marked (SM) those that did not report significant changes detected by maSigPro (Nueda et al., 2014) over time (FDR ≥ 0.05). On the contrary, differentially marked (DM) genes reported significant changes in a given mark’s profile over time (FDR < 0.05). To ensure a multiple testing correction procedure consistent among the nine marks and also with respect to gene expression, maSigPro was run, as described for gene expression (default parameters, replicates handled internally), on the initial set of 12,248 genes, which also included unmarked genes.

The next branch of classification (Fig. 4a) was applied only to the set of differentially marked genes that are also differentially expressed. To ensure consistent results among histone marks, the following multiple testing correction procedures were always applied to the set of 8,030 DE genes. For each DE gene, we computed at each time-point the breadth of a given mark’s signal, defined as the fraction of the gene’s size (from the first annotated region of interest until the last annotated Transcription Termination Site, TTS) covered by peaks of the mark. We refer to this vector of length 12 as the mark’s coverage vector. We next considered i) Pearson *r* correlation coefficient between the time-series expression levels and mark’s signals; ii) Pearson *r* correlation coefficient between the time-series expression levels and mark’s coverage values; iii) statistical significance of the Needleman-Wunch (NW) dynamic time warping alignment between the time-series expression levels and mark’s signals (following Benjamini-Hochberg multiple testing correction; script: `p-adjust.R`). We used as input for the NW alignments (scripts: `NW.alignment.path.R, NW.bidirectional.matches.py`) the z-score profiles of expression and mark obtained after applying polynomial regression `(degree = 2)` on the original matrices (scripts: `loess.polynomial. regression.R, NW.generate.input.matrix.sh`). This procedure was applied to remove the noise due to occasional fluctuations in signal over time. A permutation *p* value for each gene was computed (`script: NW.pvalue.permutation.test.py`), based on a 100,000-permutations scheme (script: `NW.alignment.permutations.R`). To classify a gene as positively correlated, we required at least two of the following conditions: i) Pearson *r* correlation coefficient between the time-series expression levels and mark’s signals ≥ 0.60 and FDR < 0.05; ii) Pearson *r* correlation coefficient between the time-series expression levels and mark’s coverage values ≥ 0.60 and FDR < 0.05; iii) NW alignment between the time-series expression levels and mark’s signals with FDR < 0.05. For negatively correlated genes, we required at least two of the following conditions: i) Pearson *r* correlation coefficient between the time-series expression levels and mark’s signals ≤ −0.60 and FDR < 0.05; ii) Pearson *r* correlation coefficient between the time-series expression levels and mark’s coverage values ≤ −0.60 and FDR < 0.05; iii) NW alignment between the time-series expression levels and mark’s signals with FDR ≥ 0.05. Genes that did not meet these requirements were classified as uncorrelated. The same decision-tree classification was performed independently for each of the nine histone marks, to ensure comparable results among all modifications (script: `define.6.groups.R`).

#### Clustering analysis

We considered all 45 combinations between the 9 histone marks and the 5 decision-tree labels described in the previous section. For instance, one combination may be “stably marked + H3K4me3”, and another combination may be “positively correlated + H3K27ac”. To test the co-occurrence of this pair of combinations, we retrieved the set of DE genes that are labelled “stably marked” for H3K4me3, and the set of DE genes that are labelled “positively correlated” for H3K27ac. The significant overlap between these two sets of genes was tested by the hypergeometric distribution (R function `phyper()`). We repeated this procedure for all possible pairs of combinations. We next clustered the *p* values obtained after applying the Benjamini-Hochberg False Discovery Rate (FDR) multiple testing correction. Hierarchical clustering was performed with the ComplexHeatmap (Gu et al., 2016) R package (clustering distance = *Manhattan*, clustering method = *Ward.D2*). Cluster correspondence analysis (van de Velden et al., 2017) of the 45 categorical variables (combinations of histone marks and decision-tree labels) across the 8,030 selected genes was performed with the R package clustrd (Markos et al., 2019). To select the optimal number of clusters and dimensions, we first run the function `tuneclus()` with the following parameters: `nclusrange = 3:10, ndimrange = 2:9, method = “clusCA”, nstart = 100, seed = 1234`. This indicated that the optimal number of dimensions and clusters was two and three, respectively. We then obtained the three clusters of genes running the function `clusmca` with the following parameters: `nclus = 3, ndim = 2, method = “clusCA”, nstart = 100, smartStart = NULL, gamma = TRUE, seed = 1234`. We obtained the same clusters of genes when running the function clusmca with the following parameters: `nclus = 3, ndim = 3, method = “MCAk”, alphak = 0.5, nstart = 100, smartStart = NULL, gamma = TRUE, seed = 1234)`. This allowed us to explore the clustering of genes also in the third dimension (Figs. 4c, S4a).

#### Gene Ontology enrichment analysis

We used the R package GOstats (Falcon and Gentleman, 2007) to identify Gene Ontology (GO) terms related to biological processes (BP) and cellular compartments (CC). We set a *p* value threshold of 0.01 to identify significantly enriched terms. For the GO enrichment analysis on the genes contributing to Principal Components (PC) 1 and 2 (described in Results, section *Gene expression recapitulates transdifferentiation more precisely than chromatin*; Fig. 1c, Table S2), we used the function `get_pca_var()` from the R package factoextra (https://CRAN.R-project.org/package=factoextra) to extract the 10% genes (n = 1,066) with the highest contribution to each of the two first principal components. The union of these two sets of genes was used as background for the GO enrichment analysis. We used REVIGO (Supek et al., 2011; http://revigo.irb.hr/) to summarize the lists of enriched GO terms. For the GO enrichment analysis on the up-regulated genes that belong to the three chromatin clusters (described in Results, section *Chromatin marking is associated with expression specifically at the time of gene activation*), we provided as background the set of 2,103 up-regulated genes. In this case, we used REVIGO and the R package ggplot2 (Wickham H, 2009) to compute and visualize, respectively, maps of the identified GO terms based on their frequency, −*log*_10_ *p* value, uniqueness and dispensability. Only children terms with dispensability < 0.5 are shown.

#### Analysis of ENCODE RNA-seq and ChIP-seq data

To investigate differences in gene expression levels and chromatin marking among the three clusters of DE genes in other biological models, we obtained RNA-seq data and ChIP-seq data for histone marks generated by the ENCODE Project (Davis et al., 2018; Dunham et al., 2012; https://www.encodeproject.org/). Besides B cells and CD14-positive monocytes, which are biologically more similar to pre-B cells and macrophages, respectively, we selected five cancer cell lines (K562, HepG2, GM12878, MCF-7, A549) that are comprehensively characterized by ENCODE ChIP-seq data for the nine histone marks that we have profiled in our study. To assess differences in gene expression levels between the three clusters of DE genes, we obtained gene expression quantifications (with respect to Gencode v24) from polyA+ RNA-seq experiments (accession date: 10/06/2019). We computed, for each gene, the average TPM values between two biological replicates. The list of experiments and datasets’ accession IDs used for this analysis is reported in Table S5.

To assess differences in chromatin marking, we obtained ChIP-seq data available for the nine histone marks profiled in our study. (Assay title: Histone ChIP-seq; Genome assembly: GRCh38; Output type: replicated peaks or stable peaks; Accession date: 10/06/2019). The list of experiments and datasets’ accession IDs used for this analysis is available in Table S6. In all cases, we excluded experiments associated with AUDIT errors. In case of multiple experiments on the same target and cell type, the experiment associated with the lowest number of AUDIT terms was selected. The scripts used to retrieve and filter the ENCODE experiments are: `download.metadata.sh, parse.metadata.audit.categories. py, retrieve.encode.identifiers.sh, parse.list.identifiers.sh`.

For each experiment and cell type, we computed the proportion of genes with at least one peak called over the gene body (H3K36me3, H4K20me1) or in the promoter region (TSS ±2 Kb for all other marks; script: `intersect.peaks.regions.sh`). In the presence of multiple TSSs annotated for the same gene, multiple regions were considered. This is consistent with the analyses described in section *ChIP-seq data processing and analysis*.

#### Analysis of temporal dynamics

For this analysis we first identified, within the set of 2,103 up-regulated genes, 257 with expression at 0 hours p.i. < 1 TPM. These genes were, therefore, specifically activated during transdifferentiation. Expression and chromatin profiles of each of the considered genes were re-scaled to range 0-100 (script: `rescale.R`): in this way, the minimum and maximum expression level or chromatin signal over the 12 time-points were set to 0% and 100% of up-regulation, respectively. We next considered, for each gene, pairs of consecutive time-points along transdifferentiation (e.g. 0h and 3h; 3h and 6h; 6h and 9h; etc.), and recorded the first time-point at which the expression / chromatin profile crossed (≥) 25%, 50%, 75% and 100% degree of up-regulation (Fig. S5b). This “crossing” step implies that, in a pair of consecutive time-points, the signal corresponding to the first time-point is, for instance, < 25%, and the signal corresponding to the second time-point is, for instance, ≥ 25%. This assessment is performed for each of the four degrees of up-regulation. To ensure monotonic increases consistently across all histone marks, we excluded genes for which this “crossing” step could not be observed for all four degrees of up-regulation in a given mark’s time-series profile. This explains the different numbers of genes, among marks, reported in Figs. 6a and S5e. For a given gene and for each of the four degrees of up-regulation, the recorded time-points (tp) for expression and chromatin profiles were compared, and a label was assigned depending on whether the up-regulation of chromatin signal anticipated (*tp_mark_* < *tp_expression_*), co-occurred (*tp_mark_* = *t_pexpression_*) or followed (*tp_mark_* > *tp_expression_*) the up-regulation of gene expression. We analogously compared the up-regulation between pairs of histone marks (Figs. 6c, S5d). In this case, we analyzed whether the up-regulation of histone mark’s signal on row *i* anticipated (*tp_i_* < *tp_j_*) or co-occurred with (*tp_i_* = *tp_j_*) the up-regulation of histone mark’s signal on column *j*. To assess whether the specific order of up-regulation in expression levels and chromatin signals depended on the initial level of expression of the genes, these analyses were repeated on a set of 629 up-regulated genes with expression at 0 hours p.i. > 25 TPM.

### QUANTIFICATION AND STATISTICAL ANALYSIS

Details regarding statistical tests, significance assessment, dispersion and precision measures are reported both in the section *Method details* and in the figures’ legends. All statistical analyses were performed using the R language for statistical computation and graphics (Team R. C., 2017; http://www.R-project.org/). In all cases, the multiple testing correction procedure was performed by applying the Benjamini-Hochberg False Discovery Rate (FDR; Benjamini and Hochberg, 1995). Wilcoxon rank-sum tests were performed with the `wiicox.test()` R function in a two-sided manner.

When not specified, plots were made using the R package ggplot2 (Wickham H, 2009). All box plots depict the first and third quartiles as the lower and upper bounds of the box, with a band inside the box showing the median value and whiskers representing 1.5x the interquartile range. All scripts used in the analyses are publicly available (see the *Data and Code Availability* statement).

## Supplemental Figures Legends

**Supplemental Figure S1: Characterization of gene expression and histone modifications’ profiles during transdifferentiation —** See also Fig. 1, Tables S1-2. **a**: Flow-cytometry plots assessing expression of CD19 and Mac-1 antigens at the 12 time-points monitored during transdifferentiation. **b**: Classification of time-series expression profiles. We selected a set of 12,248 protein-coding genes, which comprises 1,552 not expressed genes (0 TPM in all time-points and biological replicates) and 10,696 expressed genes (≥ 5 TPM in at least one time-point, and in both biological replicates). Within the set of expressed genes, we distinguished between genes with a stable expression profile throughout transdifferentiation (stably expressed; maSigPro FDR ≥ 0.05; n = 2,666), and genes showing significant changes in gene expression over time (differentially expressed or DE; maSigPro FDR < 0.05; n = 8,030). DE genes were further characterized into bending (1,409), down-regulated (4,016), peaking (502) and up-regulated (2,103) genes. Examples of genes belonging to the six types of expression profiles are provided. Gene expression values are reported in log_2_ (TPM + 1). **c**: Average pile-up signal over the gene body ± 5 Kb (H3K36me3 and H4K20me1), or promoter regions ± 5 Kb from the Transcription Start Site (TSS; all other marks), computed at each of the 12 time-points. The vertical dashed lines mark the selected region of ± 2 Kb around the TSS. **d**: Proportion of genes contributing to the first two principal components (PC1 and PC2) of the joint PCA on expression and chromatin marks (Fig. 1c), that are classified as bending, down-regulated, peaking, up-regulated or stably expressed.

**Supplemental Figure S2: The correlation between chromatin marking and gene expression over time is lower than the one reported in steady-state conditions**. — See also Fig. 1. **a**: Steady-state correlations between expression levels (x-axis) and H3K4me3 signals (y-axis) computed on the set of 12,248 genes (silent genes: dark gray; stably expressed genes: gray; DE genes: light gray). Upper panel: Pearson *r* between expression levels and H3K4me3 signals at paired time-points (0, 24, 48, 72, 120 and 168 hours). The magnitude of the correlation is reported on the top of each scatterplot. The linear regression line is depicted in brown. Lower panel: analogous representation after randomly shuffling the H3K4me3 signals among time-points. As a result, we computed the expression *vs*. chromatin correlation between unpaired time-points (0h - 24h; 24h - 120h; 48h - 168h; 72h - 0h; 120h - 72h; 168h - 48h). Steady-state correlations computed on the whole set of genes are large despite the randomization of the data. **b**: Steady-states (dots) and time-course (violin and box plots) correlation values between expression levels and chromatin signals (analogous to Fig. 1d) computed after randomly permuting the genes’ signals of a given mark among time-points. In all cases we report the Pearson *r* values averaged over 1,000 permutations. For steady-states correlations, the median Pearson *r* values across time-points are: H3K27ac: 0.63; H3K9ac: 0.68; H4K20me1: 0.54; H3K36me3: 0.69; H3K4me3: 0.69; H3K4me1: 0.50; H3K4me2: 0.59; H3K9me3: −0.08; H3K27me3: −0.16. For time-course correlations, the median Pearson *r* values across genes are 0 (|r| < 0.001) for all marks. **c**: Steady-states (dots) and time-course (violin and box plots) correlation values between expression levels and chromatin signals (analogous to Fig. 1d), computed after removing stably expressed and silent genes (i.e. only on the set of differentially expressed genes). The median steady-state Pearson *r* values for each mark are: H3K27ac: 0.53; H3K9ac: 0.58; H4K20me1: 0.56; H3K36me3: 0.60; H3K4me3: 0.51; H3K4me1: 0.17; H3K4me2: 0.26; H3K9me3: −0.08; H3K27me3: −0.23. The median time-course Pearson *r* values for each mark are: H3K27ac: 0.53; H3K9ac: 0.55; H4K20me1: 0.55; H3K36me3: 0.53; H3K4me3: 0.38; H3K4me1: 0.14; H3K4me2: 0.14; H3K9me3: 0.16; H3K27me3: −0.04. Silent genes contribute substantially to the steady state correlations, and partially contribute to the differences observed in Fig. 1d between steady-state and time-course correlations, since the latter cannot be computed for silent genes (see Methods).

**Supplemental Figure S3: Changes in chromatin marking over time can be uncoupled from changes in gene expression** — See also Figs. 2–3, Tables S3-4. **a**: Log likelihood values for HMM models with increasing number of states (between 2 and 20). **b**: Frequency of the five states observed along the twelve time-points of transdifferentiation in the HMM-sequence profiles of the sets of silent, stably expressed and differentially expressed (DE) genes. **c**: Distributions of genes’ fold-change (FC: difference between maximum and minimum signals along transdifferentiation) for each histone mark. Differences in FC among sets of silent, stably expressed and DE genes were statistically assessed with Wilcoxon Rank-Sum test (two-sided). The magnitude of chromatin changes observed in stably expressed and silent genes is, in some cases, comparable to (H4K20me1 and H3K36me3 for silent; H3K27me3 and H3K9me3 for stably expressed), or even larger (H3K4me1 and H3K4me2 for silent) than the one observed for DE genes. **d**: Trajectories of transdifferentiation derived from a Principal Component Analysis performed jointly on expression and each histone mark’s time-series profiles of DE genes, distinguishing between differentially marked (left) and stably marked (right) genes. Across all histone marks, transdifferentiation trends are clearer using the former set of genes, suggesting that the different resolution of PCA trends initially observed (Fig. 1c) may be explained by the different amount of changes observed, over time, across histone marks. Unexpectedly, H3K4me1-, H3K4me2- and H3K9me3-differentially marked genes show a contrasting profile for expression and chromatin modifications along PC2, but different to the pattern observed for H3K27me3.

**Supplemental Figure S4: Genes in different stages of activation are associated with specific chromatin and gene expression patterns, and perform distinct functions** — See also Figs. 4–5, Tables S5-6. **a**: Three-dimensional representation of the combinations of labels and histone marks (analysis attributes). The color code for the labels is analogous to Fig. 4a. Histone marks are represented by numbers. **b**: Distributions of gene expression levels at 0 and 168 hours p.i., and fold-change (FC) in gene expression (168h - 0h) for up-regulated genes that belong to clusters 1-3. Differences in gene expression levels among clusters were assessed with the Wilcoxon Rank-Sum test (two-sided). **c**: Density plot reporting the time-point at which the time-series expression profiles of up-regulated genes in clusters 1-3 reach a degree of up-regulation ≥ 25%. For this analysis, the time-series expression profile of each gene was re-scaled to a 0-100% range. **d**: Multidimensional scaling-based representation of the semantic dissimilarities between non-redundant Gene Ontology Biological Process terms enriched among up-regulated genes in clusters 1-3. Each circle represents a term, with the size and the color of the circle denoting the −*log_10_ p* value and the cluster of the term, respectively. GO terms that lie close to each other are semantically more similar. **e**: Analogous representation to Fig. S4d for cellular compartments. Cluster 1 genes are associated with metabolic functions mostly performed in intracellular compartments, suggesting a more housekeeping nature of these up-regulated genes. Cluster 2 genes perform functions related to the inflammatory response and to the cell membrane and projections, and are thus more likely to be involved in the transition from pre-B cells to macrophages. Cluster 3 genes are associated with macrophage-specific functions. **f**: Analysis of ENCODE RNA-seq data available for five cancer cell lines (MCF7, HepG2, A549, GM12878, K562), and two primary cell types (B cells and CD14+ monocytes) that are biologically similar to the cell types present at the beginning (pre-B) and at the end (macrophages), respectively, of our transdifferentiation model. Distributions of gene expression levels for up-regulated genes that belong to clusters 1-3. Differences in gene expression levels among clusters were assessed using Wilcoxon Rank-Sum test (two-sided). **g**: Analysis of ENCODE ChIP-seq data for the nine histone marks we have monitored along transdifferentiation in five cancer cell lines (MCF7, HepG2, A549, GM12878, K562) and two primary cell types (B cells and CD14+ monocytes). Proportions (%) of marked genes at gene body (H3K36me3, H4K20me1) and promoter regions (all other marks) among up-regulated genes in clusters 1-3. **h**: Percent stacked bar plot depicting the proportion of bending, down-regulated, peaking and up-regulated genes that belong to the three clusters.

**Supplemental Figure S5: The up-regulation of chromatin marks and gene expression follows a precise order only during the initial stage of gene activation** — See also Fig. 6. **a**: Alluvial plot describing the HMM time-series profiles for the 257 (upper panel) and 629 (lower panel) up-regulated genes that are not expressed (< 1 TPM) and expressed (> 25 TPM), respectively, at 0 hours p.i. **b**: Graphical representation of cases in which the up-regulation of chromatin signal anticipates (left), co-occurs with (middle), or follows (right) the up-regulation of gene expression. The expression and histone marks’ time-series profiles of each gene were re-scaled to a 0-100% range prior to the analysis. We considered four degrees of up-regulation (25%, 50%, 75% and 100%) and computed, for each gene and histone mark, the time-point at which the expression and chromatin re-scaled values reach each of the four degrees of up-regulation. Here we depict a representation for the degree of up-regulation of 25%. **c**: Lag (hours) between 50%, 75% and 100% up-regulation in histone marks’ signal and expression level for the 257 up-regulated genes not expressed at 0 hours p.i. Negative lags correspond to changes in chromatin marks anticipating changes in gene expression; positive lags correspond to changes in chromatin marks following changes in gene expression. **d**: Analogous representation to Fig. 6c for co-occurring changes between pairs of histone marks in genes that are either silent (upper panel) or expressed (lower panel) at 0 hours. For genes specifically activated during transdifferentiation (upper panel), the amount of co-occurring changes increases towards the end of the up-regulation process. **e**: Analogous representation to Fig. 6a for the 629 up-regulated genes expressed at 0 hours p.i.

**Supplemental Figure S6: Chromatin marking cannot be fully recapitulated by gene expression** — See also Figs. 1, S1. **a**: Proportion of genes contributing to the first two principal components (PC1 and PC2) of the joint PCA on expression and chromatin marks in Fig. 1c, that belong to the three clusters of DE genes (clusters 1-3) or that are stably expressed and differentially marked (“DM only”). While genes contributing to the transition from pre-B cells to macrophages (pc1-contributing genes, Fig. 1c, Fig. S1d) show the canonical correlation with chromatin changes (cluster 2), a considerable fraction of genes involved in the intermediate stages of transdifferentiation (pc2-contributing genes) display expression and chromatin changes uncoupled from one another (cluster 1, or stably expressed and differentially marked - “DM only”). This further supports the hypothesis that chromatin changes are involved in a transient de-differentiation from pre-B cells into an intermediate state, and re-differentiation into macrophages. **b**: Among the set of stably expressed and differentially marked genes contributing to PC2 (“DM only” genes in Fig. S6a), number of genes with variable chromatin profiles for increasing numbers of histone marks. For instance, 79 genes present changes in three histone modifications along transdifferentiation. **c**: Example of a stably expressed gene (*TALDO1*) contributing to PC2 in the PCA in Fig. 1c, and showing significant changes in some chromatin profiles along transdifferentiation. Expression and chromatin tracks from one biological replicate are displayed, as well as normalized line plots averaging the signal from the two replicates. Profiles of HMM states are shown at the bottom.

## Supplemental Tables Legends

**Supplemental Table S1: Numbers of unmarked and marked genes within the sets of 1,552 silent and 10,696 expressed genes —** See also Figs. 1, S1. For a given histone mark, unmarked genes have no peaks called at any time-point in the region of interest, while marked genes have peaks called in the region of interest in at least one time-point (see Methods).

**Supplemental Table S2: GO terms significantly enriched among genes contributing to Principal Components 1 and 2** — See also Figs. 1, S1. The list of terms refers to Biological Processes.

**Supplemental Table S3: Decision-tree labelling of differentially expressed genes** — See also Figs. 3–4, S3. Left side: numbers of unmarked and marked genes within the set of DE genes. Marked genes are further separated into genes that are either stably or differentially marked (i.e. have stable or variable chromatin profiles during transdifferentiation). The percentages refer to the total number of DE genes (n = 8,030). Right side: within the set of differentially marked genes, we distinguish between genes that are positively correlated, uncorrelated or negatively correlated with gene expression over time (see Methods). The percentages in this case are computed with respect to the number of differentially marked genes found for each histone modification.

**Supplemental Table S4: Absent, stable and differential chromatin marking over time among silent and stably expressed genes** — See also Figs. 3, S3. Numbers of unmarked and marked genes within the sets of 1,552 silent and 2,666 stably expressed genes. Marked genes are further separated into genes that are either stably or differentially marked.

**Supplemental Table S5: ENCODE PolyA+ RNA-seq experiments in seven cell types** — See also Fig. S4. The ENCODE accession numbers allow to uniquely identify the experiment and gene expression quantification file (tsv) on the ENCODE portal (https://www.encodeproject.org/).

**Supplemental Table S6: ENCODE histone ChIP-seq experiments in seven cell types** — See also Fig. S4. The ENCODE accession numbers allow to uniquely identify the experiment and peak call file (bigBed) on the ENCODE portal (https://www.encodeproject.org/).

**Supplemental Table S7: Catalog of 12,248 protein-coding genes analyzed in this study**. For each gene we provide the level of expression at 0 hours p.i. (average of the normalized levels from the two biological replicates), the type of expression profile (silent / stably expressed / bending / down-regulated / peaking / up-regulated) and the chromatin marking status (unmarked / stably marked / differentially marked) with respect to each of the nine histone marks. In the case of DE genes, we further specify the type of relationship with gene expression over time (positively correlated / uncorrelated / negatively correlated), as well as the corresponding chromatin cluster (1: stable / uncorrelated marking; 2: positively correlated marking; 3: absence of marking).

## Notes

### Competing Interest Statement

The authors have declared no competing interest.

