## Supplementary material for "Dynamics of gene expression and chromatin marking during cell state transition": SI.merged.pdf

**Supplemental Figure S1**

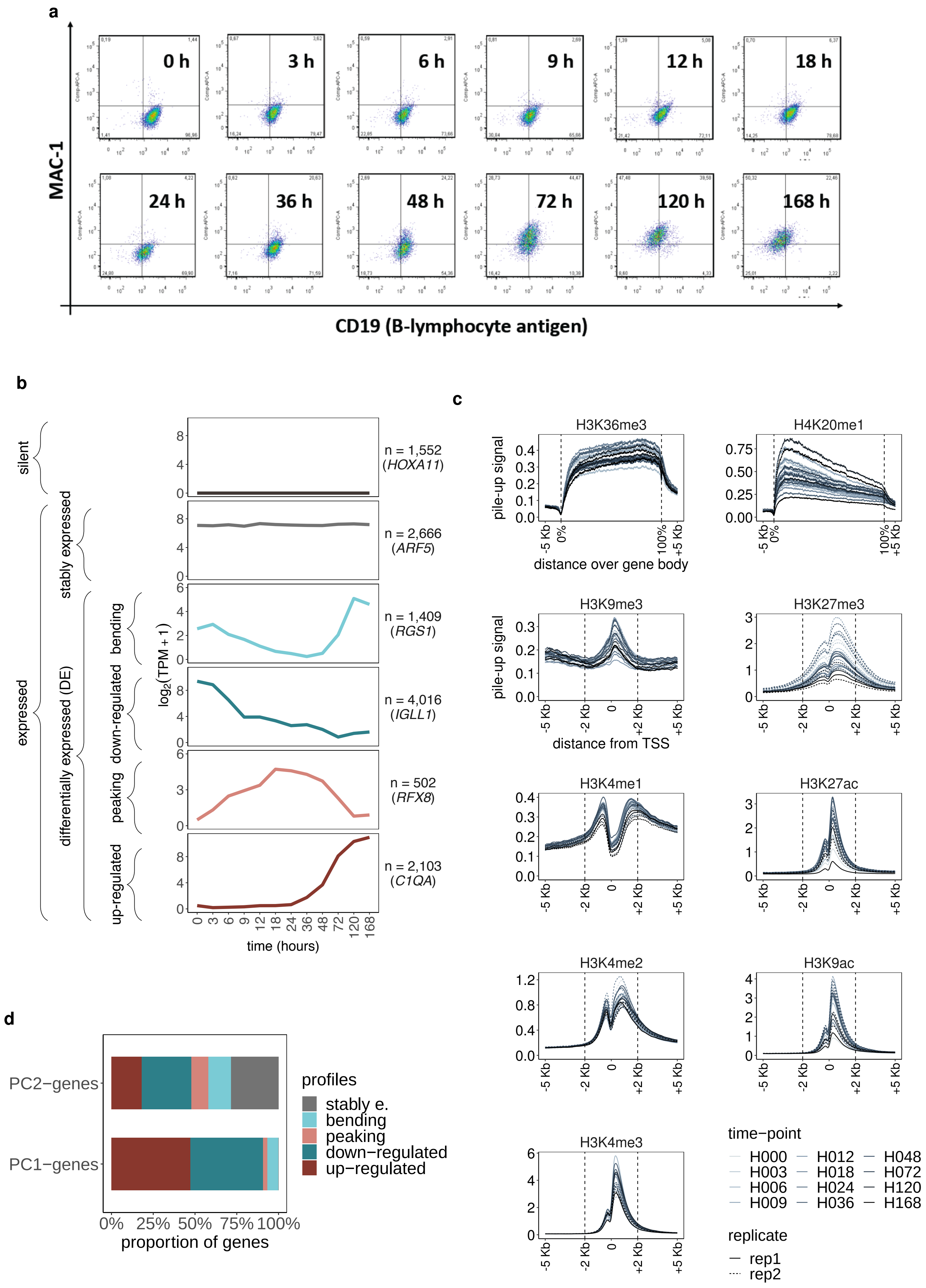

**Supplemental Figure S1: Characterization of gene expression and histone modifications' profiles during transdifferentiation** — See also Fig. 1, Tables S1-2. **a**: Flow-cytometry plots assessing expression of CD19 and Mac-1 antigens at the 12 time-points monitored during transdifferentiation. **b**: Classification of time-series expression profiles. We selected a set of 12,248 protein-coding genes, which comprises 1,552 not expressed genes (0 TPM in all time-points and biological replicates) and 10,696 expressed genes ($\geq$ 5 TPM in at least one time-point, and in both biological replicates). Within the set of expressed genes, we distinguished between genes with a stable expression profile throughout transdifferentiation (stably expressed; maSigPro FDR $\geq$ 0.05; n = 2,666), and genes showing significant changes in gene expression over time (differentially expressed or DE; maSigPro FDR $<$ 0.05; n = 8,030). DE genes were further characterized into bending (1,409), down-regulated (4,016), peaking (502) and up-regulated (2,103) genes. Examples of genes belonging to the six types of expression profiles are provided. Gene expression values are reported in $\log_2$ (TPM + 1). **c**: Average pile-up signal over the gene body $\pm$ 5 Kb (H3K36me3 and H4K20me1), or promoter regions $\pm$ 5 Kb from the Transcription Start Site (TSS; all other marks), computed at each of the 12 time-points. The vertical dashed lines mark the selected region of $\pm$ 2 Kb around the TSS. **d**: Proportion of genes contributing to the first two principal components (PC1 and PC2) of the joint PCA on expression and chromatin marks (Fig. 1c), that are classified as bending, down-regulated, peaking, up-regulated or stably expressed.

**Supplemental Figure S2**

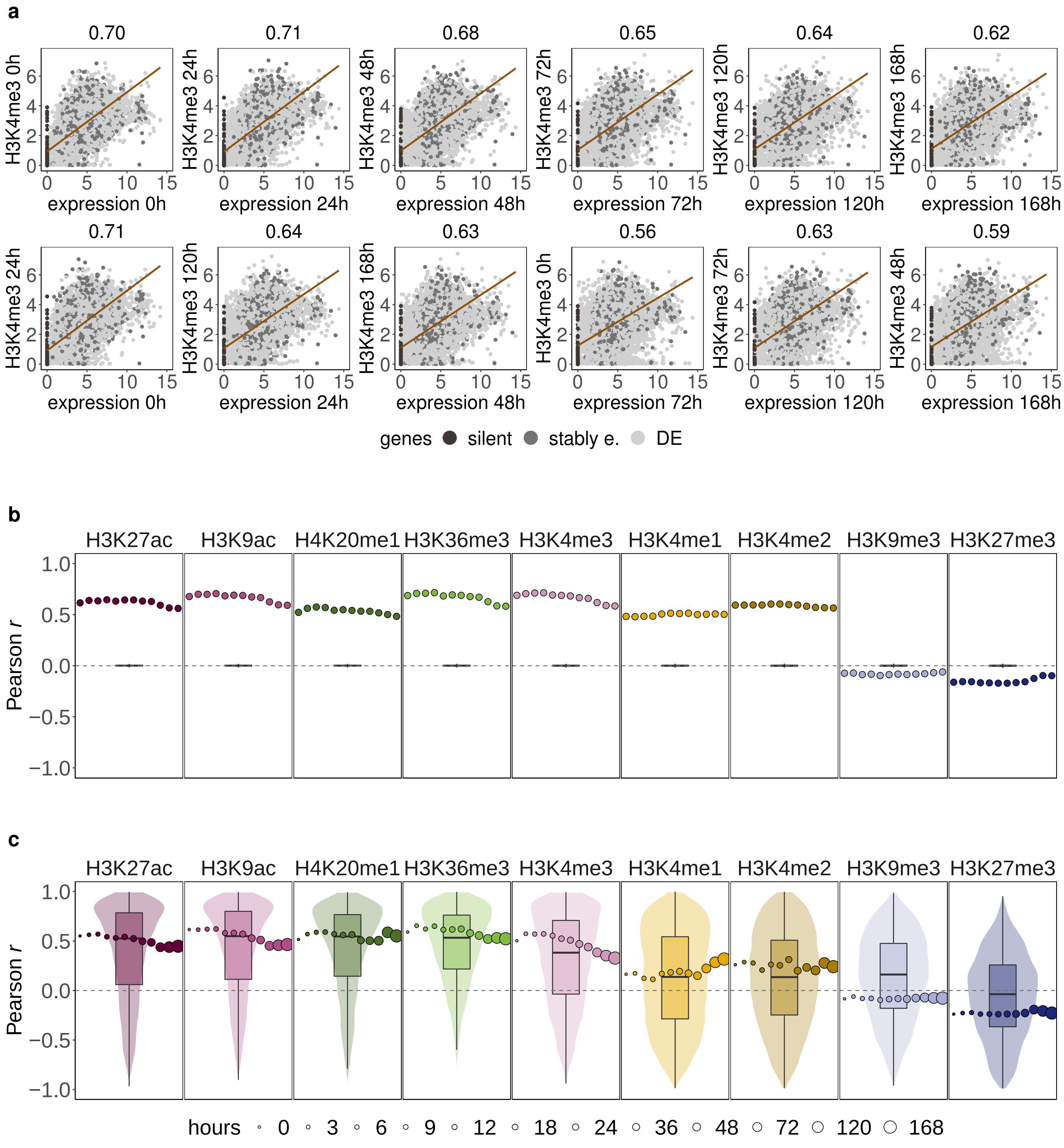

**Supplemental Figure S2: The correlation between chromatin marking and gene expression over time is lower than the one reported in steady-state conditions.** — See also Fig. 1. **a**: Steady-state correlations between expression levels (x-axis) and H3K4me3 signals (y-axis) computed on the set of 12,248 genes (silent genes: dark gray; stably expressed genes: gray; DE genes: light gray). Upper panel: Pearson $r$ between expression levels and H3K4me3 signals at paired time-points (0, 24, 48, 72, 120 and 168 hours). The magnitude of the correlation is reported on the top of each scatterplot. The linear regression line is depicted in brown. Lower panel: analogous representation after randomly shuffling the H3K4me3 signals among time-points. As a result, we computed the expression *vs.* chromatin correlation between unpaired time-points (0h - 24h; 24h - 120h; 48h - 168h; 72h - 0h; 120h - 72h; 168h - 48h). Steady-state correlations computed on the whole set of genes are large despite the randomization of the data. **b**: Steady-states (dots) and time-course (violin and box plots) correlation values between expression levels and chromatin signals (analogous to Fig. 1d) computed after randomly permuting the genes' signals of a given mark among time-points. In all cases we report the Pearson $r$ values averaged over 1,000 permutations. For steady-states correlations, the median Pearson $r$ values across time-points are: H3K27ac: 0.63; H3K9ac: 0.68; H4K20me1: 0.54; H3K36me3: 0.69; H3K4me3: 0.69; H3K4me1: 0.50; H3K4me2: 0.59; H3K9me3: -0.08; H3K27me3: -0.16. For time-course correlations, the median Pearson $r$ values across genes are 0 ($|r| < 0.001$) for all marks. **c**: Steady-states (dots) and time-course (violin and box plots) correlation values between expression levels and chromatin signals (analogous to Fig. 1d), computed after removing stably expressed and silent genes (i.e. only on the set of differentially expressed genes). The median steady-state Pearson $r$ values for each mark are: H3K27ac: 0.53; H3K9ac: 0.58; H4K20me1: 0.56; H3K36me3: 0.60; H3K4me3: 0.51; H3K4me1: 0.17; H3K4me2: 0.26; H3K9me3: -0.08; H3K27me3: -0.23. The median time-course Pearson $r$ values for each mark are: H3K27ac: 0.53; H3K9ac: 0.55; H4K20me1: 0.55; H3K36me3: 0.53; H3K4me3: 0.38; H3K4me1: 0.14; H3K4me2: 0.14; H3K9me3: 0.16; H3K27me3: -0.04. Silent genes contribute substantially to the steady state correlations, and partially contribute to the differences observed in Fig. 1d between steady-state and time-course correlations, since the latter cannot be computed for silent genes (see Methods).

### Supplemental Figure S3

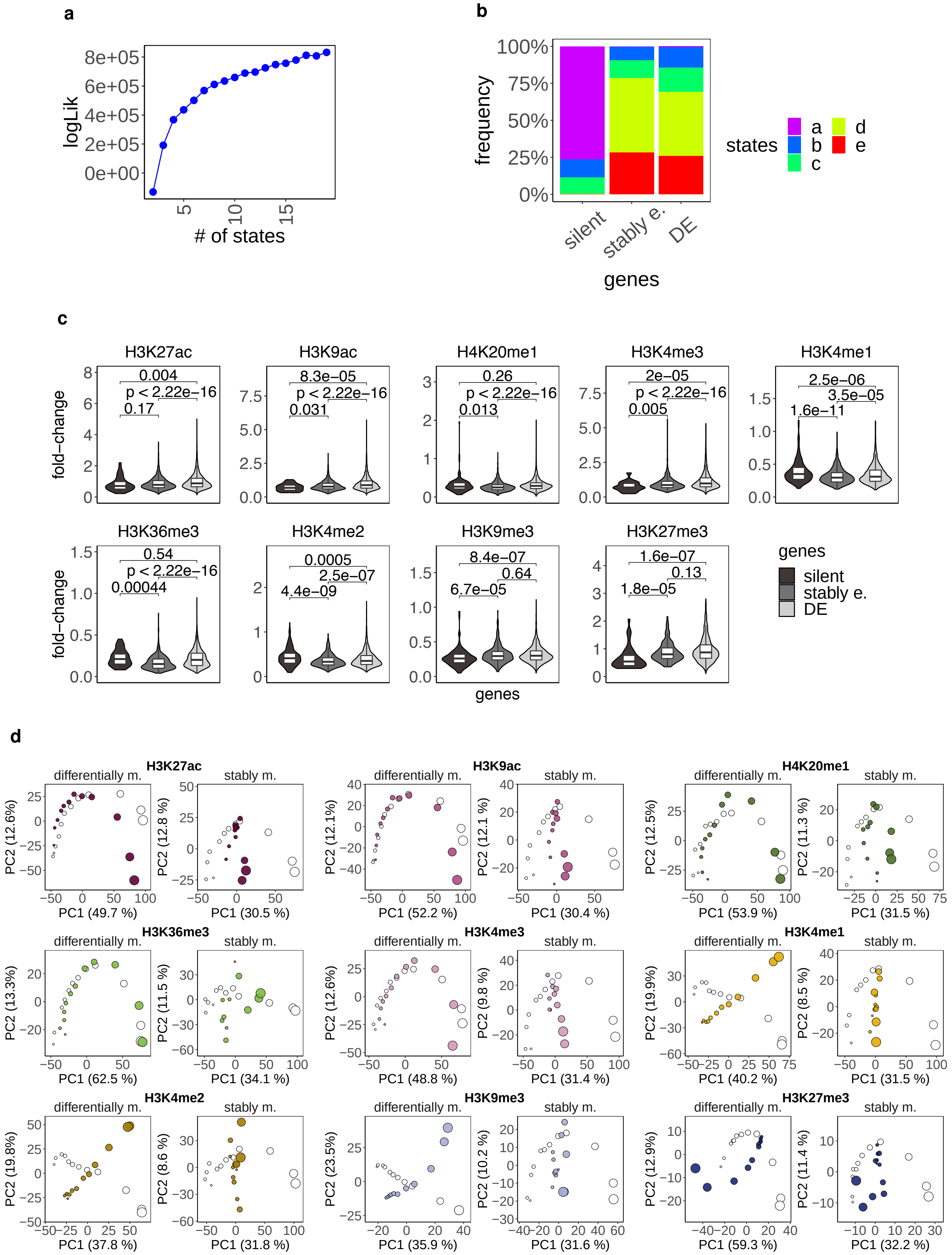

**Supplemental Figure S3: Changes in chromatin marking over time can be uncoupled from changes in gene expression** — See also Figs. 2-3, Tables S3-4. **a**: Log likelihood values for HMM models with increasing number of states (between 2 and 20). **b**: Frequency of the five states observed along the twelve time-points of transdifferentiation in the HMM-sequence profiles of the sets of silent, stably expressed and differentially expressed (DE) genes. **c**: Distributions of genes' fold-change (FC: difference between maximum and minimum signals along transdifferentiation) for each histone mark. Differences in FC among sets of silent, stably expressed and DE genes were statistically assessed with Wilcoxon Rank-Sum test (two-sided). The magnitude of chromatin changes observed in stably expressed and silent genes is, in some cases, comparable to (H4K20me1 and H3K36me3 for silent; H3K27me3 and H3K9me3 for stably expressed), or even larger (H3K4me1 and H3K4me2 for silent) than the one observed for DE genes. **d**: Trajectories of transdifferentiation derived from a Principal Component Analysis performed jointly on expression and each histone mark's time-series profiles of DE genes, distinguishing between differentially marked (left) and stably marked (right) genes. Across all histone marks, transdifferentiation trends are clearer using the former set of genes, suggesting that the different resolution of PCA trends initially observed (Fig. 1c) may be explained by the different amount of changes observed, over time, across histone marks. Unexpectedly, H3K4me1-, H3K4me2- and H3K9me3-differentially marked genes show a contrasting profile for expression and chromatin modifications along PC2, but different to the pattern observed for H3K27me3.

### Supplemental Figure S4

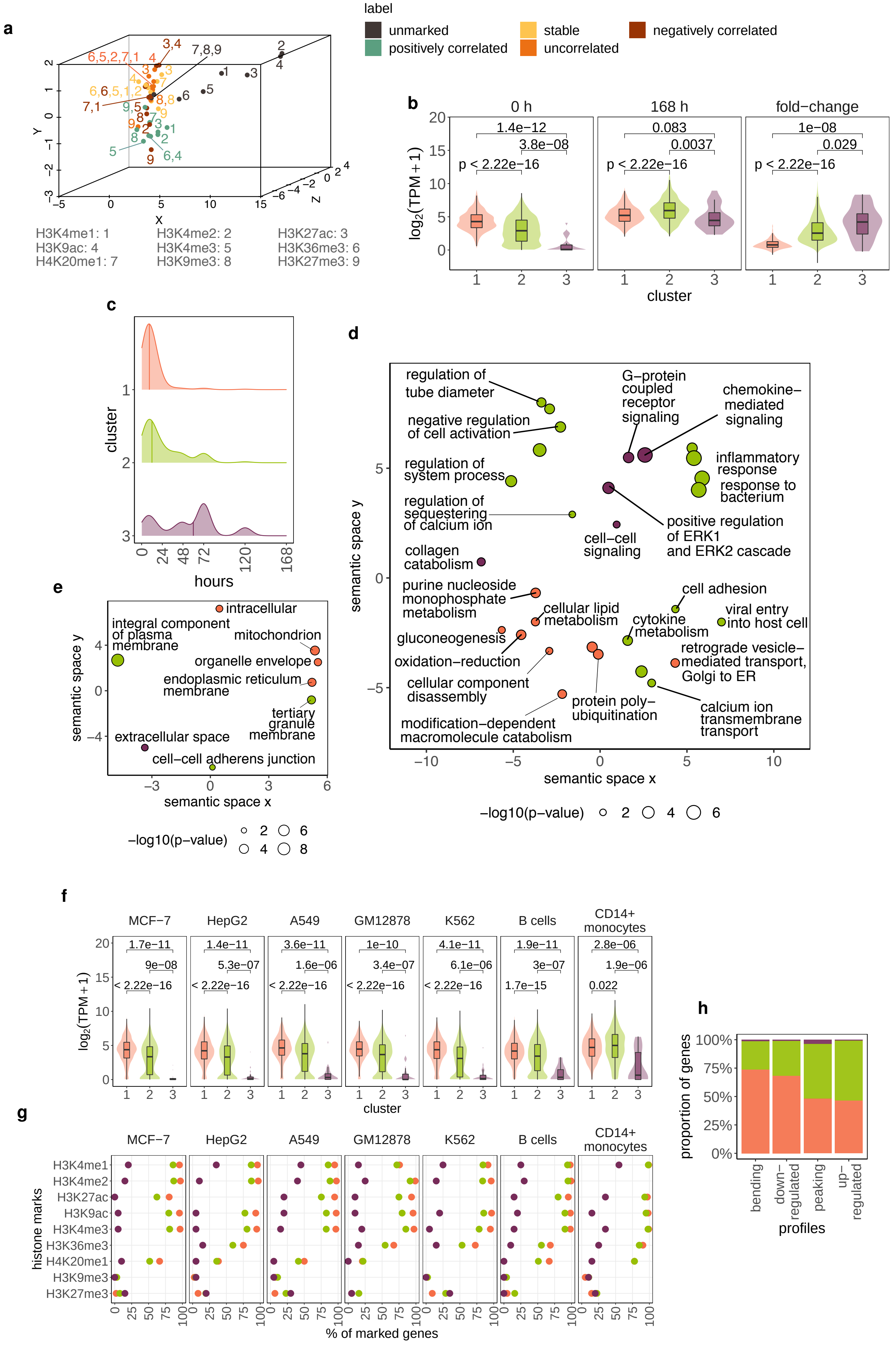

**Supplemental Figure S4: Genes in different stages of activation are associated with specific chromatin and gene expression patterns, and perform distinct functions** — See also Figs. 4-5, Tables S5-6. **a**: Three-dimensional representation of the combinations of labels and histone marks (analysis attributes). The color code for the labels is analogous to Fig. 4a. Histone marks are represented by numbers. **b**: Distributions of gene expression levels at 0 and 168 hours p.i., and fold-change (FC) in gene expression (168h - 0h) for up-regulated genes that belong to clusters 1-3. Differences in gene expression levels among clusters were assessed with the Wilcoxon Rank-Sum test (two-sided). **c**: Density plot reporting the time-point at which the time-series expression profiles of up-regulated genes in clusters 1-3 reach a degree of up-regulation $\geq$ 25%. For this analysis, the time-series expression profile of each gene was re-scaled to a 0-100% range. **d**: Multidimensional scaling-based representation of the semantic dissimilarities between non-redundant Gene Ontology Biological Process terms enriched among up-regulated genes in clusters 1-3. Each circle represents a term, with the size and the color of the circle denoting the $-log_{10}$ *p* value and the cluster of the term, respectively. GO terms that lie close to each other are semantically more similar. **e**: Analogous representation to Fig. S4d for cellular compartments. Cluster 1 genes are associated with metabolic functions mostly performed in intracellular compartments, suggesting a more housekeeping nature of these up-regulated genes. Cluster 2 genes perform functions related to the inflammatory response and to the cell membrane and projections, and are thus more likely to be involved in the transition from pre-B cells to macrophages. Cluster 3 genes are associated with macrophage-specific functions. **f**: Analysis of ENCODE RNA-seq data available for five cancer cell lines (MCF7, HepG2, A549, GM12878, K562), and two primary cell types (B cells and CD14+ monocytes) that are biologically similar to the cell types present at the beginning (pre-B) and at the end (macrophages), respectively, of our transdifferentiation model. Distributions of gene expression levels for up-regulated genes that belong to clusters 1-3. Differences in gene expression levels among clusters were assessed using Wilcoxon Rank-Sum test (two-sided). **g**: Analysis of ENCODE ChIP-seq data for the nine histone marks we have monitored along transdifferentiation in five cancer cell lines (MCF7, HepG2, A549, GM12878, K562) and two primary cell types (B cells and CD14+ monocytes). Proportions (%) of marked genes at gene body (H3K36me3, H4K20me1) and promoter regions (all other marks) among up-regulated genes in clusters 1-3. **h**: Percent stacked bar plot depicting the proportion of bending, down-regulated, peaking and up-regulated genes that belong to the three clusters.

**Supplemental Figure S5**

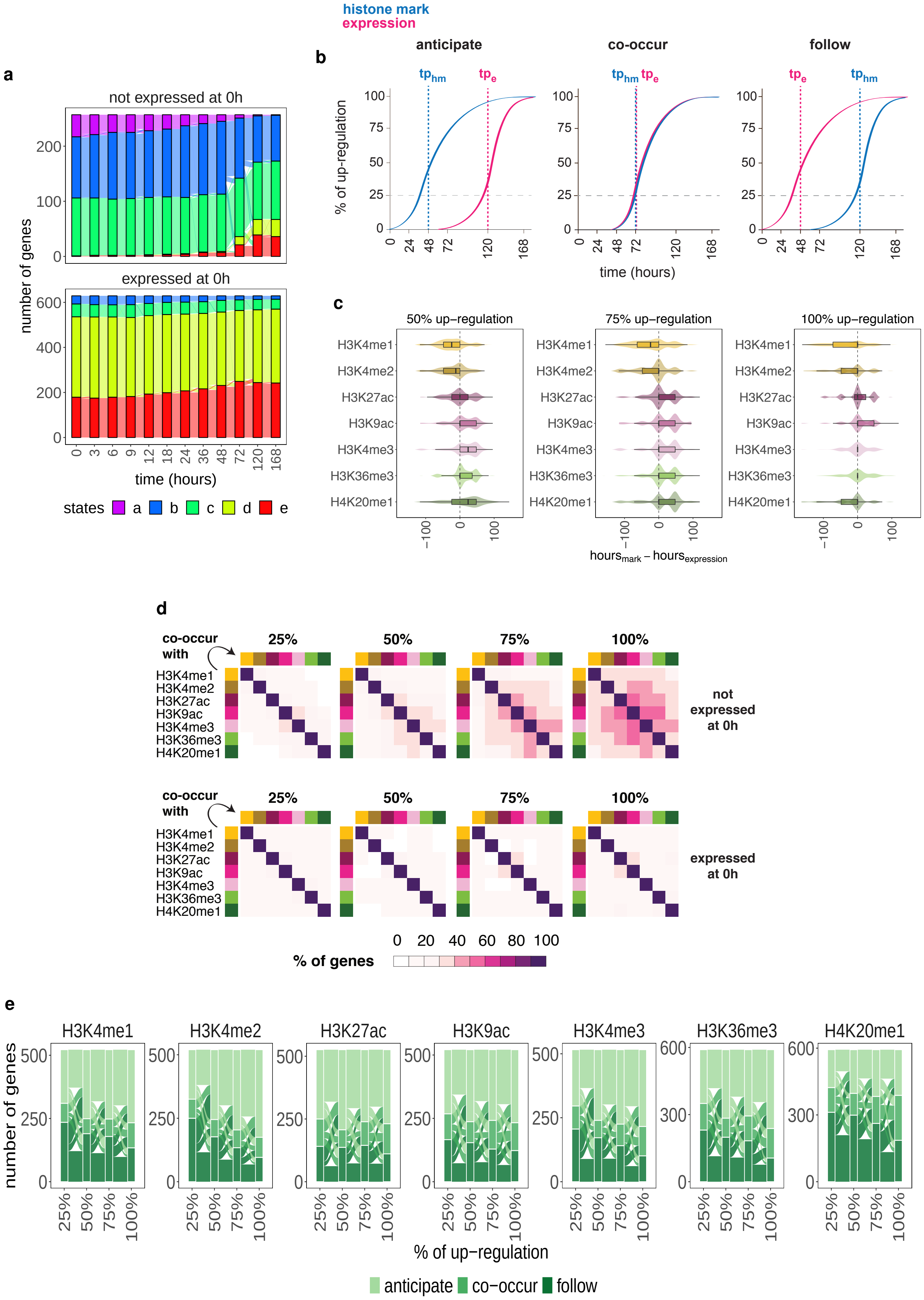

**Supplemental Figure S5: The up-regulation of chromatin marks and gene expression follows a precise order only during the initial stage of gene activation** — See also Fig. 6. **a**: Alluvial plot describing the HMM time-series profiles for the 257 (upper panel) and 629 (lower panel) up-regulated genes that are not expressed ($<$ 1 TPM) and expressed ($>$ 25 TPM), respectively, at 0 hours p.i. **b**: Graphical representation of cases in which the up-regulation of chromatin signal anticipates (left), co-occurs with (middle), or follows (right) the up-regulation of gene expression. The expression and histone marks' time-series profiles of each gene were re-scaled to a 0-100% range prior to the analysis. We considered four degrees of up-regulation (25%, 50%, 75% and 100%) and computed, for each gene and histone mark, the time-point at which the expression and chromatin re-scaled values reach each of the four degrees of up-regulation. Here we depict a representation for the degree of up-regulation of 25%. **c**: Lag (hours) between 50%, 75% and 100% up-regulation in histone marks' signal and expression level for the 257 up-regulated genes not expressed at 0 hours p.i. Negative lags correspond to changes in chromatin marks anticipating changes in gene expression; positive lags correspond to changes in chromatin marks following changes in gene expression. **d**: Analogous representation to Fig. 6c for co-occurring changes between pairs of histone marks in genes that are either silent (upper panel) or expressed (lower panel) at 0 hours. For genes specifically activated during transdifferentiation (upper panel), the amount of co-occurring changes increases towards the end of the up-regulation process. **e**: Analogous representation to Fig. 6a for the 629 up-regulated genes expressed at 0 hours p.i.

**Supplemental Figure S6**

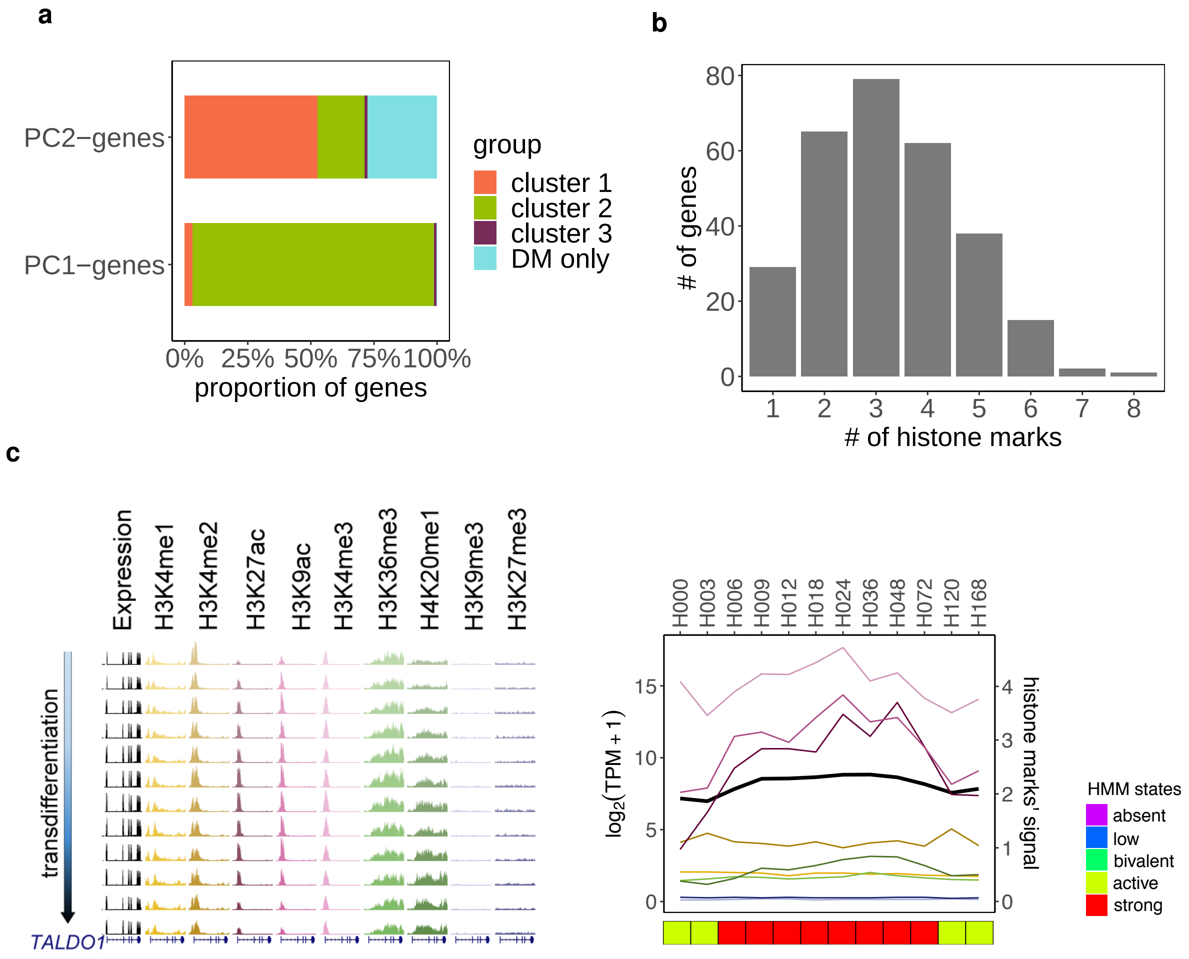

**Supplemental Figure S6: Chromatin marking cannot be fully recapitulated by gene expression** — See also Figs. 1, S1. **a**: Proportion of genes contributing to the first two principal components (PC1 and PC2) of the joint PCA on expression and chromatin marks in Fig. 1c, that belong to the three clusters of DE genes (clusters 1-3) or that are stably expressed and differentially marked (“DM only”). While genes contributing to the transition from pre-B cells to macrophages (pc1-contributing genes, Fig. 1c, Fig. S1d) show the canonical correlation with chromatin changes (cluster 2), a considerable fraction of genes involved in the intermediate stages of transdifferentiation (pc2-contributing genes) display expression and chromatin changes uncoupled from one another (cluster 1, or stably expressed and differentially marked - “DM only”). This further supports the hypothesis that chromatin changes are involved in a transient de-differentiation from pre-B cells into an intermediate state, and re-differentiation into macrophages. **b**: Among the set of stably expressed and differentially marked genes contributing to PC2 (“DM only” genes in Fig. S6a), number of genes with variable chromatin profiles for increasing numbers of histone marks. For instance, 79 genes present changes in three histone modifications along transdifferentiation. **c**: Example of a stably expressed gene (*TALDO1*) contributing to PC2 in the PCA in Fig. 1c, and showing significant changes in some chromatin profiles along transdifferentiation. Expression and chromatin tracks from one biological replicate are displayed, as well as normalized line plots averaging the signal from the two replicates. Profiles of HMM states are shown at the bottom.

**Supplemental Table S1**

| histone marks | silent | | expressed | |
|---|---|---|---|---|
| | unmarked | marked | unmarked | marked |
| H3K4me1 | 1,165 (75.1%) | 387 (24.9%) | 52 (0.5%) | 10,644 (99.5%) |
| H3K4me2 | 1,225 (78.9%) | 327 (21.1%) | 45 (0.4%) | 10,651 (99.6%) |
| H3K9ac | 1,500 (96.6%) | 52 (3.4%) | 130 (1.2%) | 10,566 (98.8%) |
| H3K27ac | 1,478 (95.2%) | 74 (4.8%) | 159 (1.5%) | 10,537 (98.5%) |
| H3K4me3 | 1,488 (95.9%) | 64 (4.1%) | 183 (1.7%) | 10,513 (98.3%) |
| H3K36me3 | 1,444 (93.0%) | 108 (7.0%) | 354 (3.3%) | 10,342 (96.7%) |
| H4K20me1 | 1,348 (86.9%) | 204 (13.1%) | 1,851 (17.3%) | 8,845 (82.7%) |
| H3K9me3 | 990 (63.8%) | 562 (36.2%) | 7,201 (67.3%) | 3,495 (32.7%) |
| H3K27me3 | 1,371 (88.3%) | 181 (11.7%) | 9,421 (88.1%) | 1,275 (11.9%) |

**Supplemental Table S1: Numbers of unmarked and marked genes within the sets of 1,552 silent and 10,696 expressed genes** — See also Figs. 1, S1. For a given histone mark, unmarked genes have no peaks called at any time-point in the region of interest, while marked genes have peaks called in the region of interest in at least one time-point (see Methods).

**Supplemental Table S3**

| histone marks | unmarked | marked | | differentially marked | | |
|---|---|---|---|---|---|---|
| | | stably | differentially | positively c. | uncorrelated | negatively c. |
| H3K4me1 | 41 (0.5%) | 4,591 (57.2%) | 3,398 (42.3%) | 1,491 (43.9%) | 1,457 (42.9%) | 450 (13.2%) |
| H3K4me2 | 32 (0.4%) | 5,074 (63.2%) | 2,924 (36.4%) | 1,248 (42.7%) | 1,316 (45.0%) | 360 (12.3%) |
| H3K9ac | 107 (1.3%) | 2,996 (37.3%) | 4,927 (61.4%) | 3,239 (65.8%) | 1,548 (31.4%) | 140 (2.8%) |
| H3K27ac | 135 (1.7%) | 2,835 (35.3%) | 5,060 (63.0%) | 3,065 (60.6%) | 1,761 (34.8%) | 234 (4.6%) |
| H3K4me3 | 150 (1.9%) | 4,310 (53.7%) | 3,570 (44.4%) | 2,048 (57.4%) | 1,402 (39.3%) | 120 (3.3%) |
| H3K36me3 | 283 (3.5%) | 4,801 (59.8%) | 2,946 (36.7%) | 2,472 (83.9%) | 459 (15.6%) | 15 (0.5%) |
| H4K20me1 | 1,403 (17.5%) | 2,484 (30.9%) | 4,143 (51.6%) | 2,782 (67.2%) | 1,256 (30.3%) | 105 (2.5%) |
| H3K9me3 | 5,363 (66.8%) | 1,698 (21.1%) | 969 (12.1%) | 253 (26.1%) | 615 (63.5%) | 101 (10.4%) |
| H3K27me3 | 6,988 (87.0%) | 362 (4.5%) | 680 (8.5%) | 40 (5.9%) | 347 (51.0%) | 293 (43.1%) |

**Supplemental Table S3: Decision-tree labelling of differentially expressed genes** — See also Figs. 3-4, S3. Left side: numbers of unmarked and marked genes within the set of DE genes. Marked genes are further separated into genes that are either stably or differentially marked (i.e. have stable or variable chromatin profiles during transdifferentiation). The percentages refer to the total number of DE genes (n = 8,030). Right side: within the set of differentially marked genes, we distinguish between genes that are positively correlated, uncorrelated or negatively correlated with gene expression over time (see Methods). The percentages in this case are computed with respect to the number of differentially marked genes found for each histone modification.

**Supplemental Table S4**

| histone mark | silent | | | stably expressed | | |
|---|---|---|---|---|---|---|
| | unmarked | marked | | unmarked | marked | |
| | | stably | differentially | | stably | differentially |
| H3K4me1 | 1,165 (75.1%) | 114 (7.3%) | 273 (17.6%) | 11 (0.4%) | 1,711 (64.2%) | 944 (35.4%) |
| H3K4me2 | 1,225 (78.9%) | 91 (5.9%) | 236 (15.2%) | 13 (0.5%) | 1,904 (71.4%) | 749 (28.1%) |
| H3K9ac | 1,500 (96.6%) | 15 (1%) | 37 (2.4%) | 23 (0.9%) | 1,338 (50.2%) | 1,305 (48.9%) |
| H3K27ac | 1,478 (95.2%) | 28 (1.8%) | 46 (3%) | 24 (0.9%) | 1,197 (44.9%) | 1,445 (54.2%) |
| H3K4me3 | 1,488 (95.9%) | 30 (1.9%) | 34 (2.2%) | 33 (1.2%) | 1,741 (65.3%) | 892 (33.5%) |
| H3K36me3 | 1,444 (93%) | 78 (5%) | 30 (1.9%) | 71 (2.7%) | 2,204 (82.7%) | 391 (14.7%) |
| H4K20me1 | 1,348 (86.9%) | 88 (5.7%) | 116 (7.5%) | 448 (16.8%) | 1,221 (45.8%) | 997 (37.4%) |
| H3K9me3 | 990 (63.8%) | 445 (28.7%) | 117 (7.5%) | 1,838 (68.9%) | 558 (20.9%) | 270 (10.1%) |
| H3K27me3 | 1,371 (88.3%) | 138 (8.9%) | 43 (2.8%) | 2,433 (91.3%) | 125 (4.7%) | 108 (4.1%) |

**Supplemental Table S4: Absent, stable and differential chromatin marking over time among silent and stably expressed genes** — See also Figs. 3, S3. Numbers of unmarked and marked genes within the sets of 1,552 silent and 2,666 stably expressed genes. Marked genes are further separated into genes that are either stably or differentially marked.

**Supplemental Table S5**

| Experiment ID | Accession file ID | Replicate | Biosample term name |
|---|---|---|---|
| ENCSR000CON | ENCFF369ZNM | 1 | A549 |
| ENCSR000CON | ENCFF627QMV | 2 | A549 |
| ENCSR000CTV | ENCFF485EUP | 1 | B cell |
| ENCSR000CTV | ENCFF231GYC | 2 | B cell |
| ENCSR000CUC | ENCFF299BIL | 1 | CD14-positive monocyte |
| ENCSR000CUC | ENCFF397DFK | 2 | CD14-positive monocyte |
| ENCSR000AED | ENCFF902UYP | 1 | GM12878 |
| ENCSR000AED | ENCFF550OHK | 2 | GM12878 |
| ENCSR000CPE | ENCFF004HYK | 1 | HepG2 |
| ENCSR000CPE | ENCFF401KRE | 2 | HepG2 |
| ENCSR000CPH | ENCFF172GIN | 1 | K562 |
| ENCSR000CPH | ENCFF768TKT | 2 | K562 |
| ENCSR000CPT | ENCFF009GDJ | 1 | MCF-7 |
| ENCSR000CPT | ENCFF885LEQ | 2 | MCF-7 |

**Supplemental Table S5: ENCODE PolyA+ RNA-seq experiments in seven cell types** — See also Fig. S4. The ENCODE accession numbers allow to uniquely identify the experiment and gene expression quantification file (tsv) on the ENCODE portal (https://www.encodeproject.org/).

**Supplemental Table S6**

| Histone mark | Experiment ID | Accession file ID | Biosample term name |
|---|---|---|---|
| H3K27ac | ENCSR000AUI | ENCFF268BMM | A549 |
| H3K27me3 | ENCSR000AUK | ENCFF368SNX | A549 |
| H3K4me1 | ENCSR000AUM | ENCFF761SFV | A549 |
| H3K4me2 | ENCSR000AVI | ENCFF260MGY | A549 |
| H3K4me3 | ENCSR000DPD | ENCFF820IQP | A549 |
| H3K9ac | ENCSR000ASV | ENCFF649ABE | A549 |
| H3K9me3 | ENCSR000AUN | ENCFF900ULD | A549 |
| H4K20me1 | ENCSR000AUO | ENCFF505MWT | A549 |
| H3K27ac | ENCSR000AUP | ENCFF041HKG | B cell |
| H3K27me3 | ENCSR162DGX | ENCFF428KOX | B cell |
| H3K36me3 | ENCSR424XBP | ENCFF649XGE | B cell |
| H3K4me1 | ENCSR290YLQ | ENCFF778RHF | B cell |
| H3K4me2 | ENCSR000AUY | ENCFF615MAT | B cell |
| H3K4me3 | ENCSR878JSF | ENCFF225QYU | B cell |
| H3K9ac | ENCSR799SLA | ENCFF890NWX | B cell |
| H3K9me3 | ENCSR005WWZ | ENCFF281WGS | B cell |
| H4K20me1 | ENCSR000AVJ | ENCFF856EUL | B cell |
| H3K27ac | ENCSR000ASJ | ENCFF239LOH | CD14-positive monocyte |
| H3K27me3 | ENCSR000ASK | ENCFF930KLN | CD14-positive monocyte |
| H3K36me3 | ENCSR000ASL | ENCFF108MXF | CD14-positive monocyte |
| H3K4me1 | ENCSR000ASM | ENCFF673ZGJ | CD14-positive monocyte |
| H3K4me3 | ENCSR000ASN | ENCFF691MBD | CD14-positive monocyte |
| H3K9ac | ENCSR000ATF | ENCFF994MCP | CD14-positive monocyte |
| H3K9me3 | ENCSR000ASP | ENCFF236ADT | CD14-positive monocyte |
| H4K20me1 | ENCSR000ASQ | ENCFF887JRI | CD14-positive monocyte |
| H3K27ac | ENCSR000AKC | ENCFF690GQK | GM12878 |
| H3K27me3 | ENCSR000DRX | ENCFF103NGB | GM12878 |
| H3K36me3 | ENCSR000DRW | ENCFF144MAY | GM12878 |
| H3K4me1 | ENCSR000AKF | ENCFF378FBA | GM12878 |
| H3K4me2 | ENCSR000AKG | ENCFF514YHH | GM12878 |
| H3K4me3 | ENCSR057BWO | ENCFF296PTF | GM12878 |

| | | | |
|---|---|---|---|
| H3K9ac | ENCSR000AKH | ENCFF637GBK | GM12878 |
| H4K20me1 | ENCSR000AKI | ENCFF154MVT | GM12878 |
| H3K27me3 | ENCSR000DUE | ENCFF034QJR | HepG2 |
| H3K36me3 | ENCSR000DUD | ENCFF370NTL | HepG2 |
| H3K4me1 | ENCSR000APV | ENCFF095ZHO | HepG2 |
| H3K4me2 | ENCSR000AMC | ENCFF948LWD | HepG2 |
| H3K4me3 | ENCSR575RRX | ENCFF229PGV | HepG2 |
| H3K9ac | ENCSR000AMD | ENCFF129YID | HepG2 |
| H3K9me3 | ENCSR000ATD | ENCFF997LPG | HepG2 |
| H4K20me1 | ENCSR000AMQ | ENCFF031AYD | HepG2 |
| H3K27me3 | ENCSR000EWB | ENCFF233ODK | K562 |
| H3K36me3 | ENCSR000DWB | ENCFF514DBT | K562 |
| H3K4me1 | ENCSR000EWC | ENCFF359WWB | K562 |
| H3K4me2 | ENCSR000AKT | ENCFF168NKC | K562 |
| H3K4me3 | ENCSR668LDD | ENCFF465RJJ | K562 |
| H3K9ac | ENCSR000EVZ | ENCFF257END | K562 |
| H3K9me3 | ENCSR000APE | ENCFF361WTS | K562 |
| H3K27ac | ENCSR000EWR | ENCFF040ZCD | MCF-7 |
| H3K27me3 | ENCSR000EWP | ENCFF825FPO | MCF-7 |
| H3K4me1 | ENCSR493NBY | ENCFF158SKW | MCF-7 |
| H3K4me2 | ENCSR875KOJ | ENCFF651IUJ | MCF-7 |
| H3K4me3 | ENCSR000DWJ | ENCFF530SPD | MCF-7 |
| H3K9ac | ENCSR056UBA | ENCFF636NEF | MCF-7 |
| H3K9me3 | ENCSR000EWQ | ENCFF348ISZ | MCF-7 |
| H4K20me1 | ENCSR639RHG | ENCFF052ILJ | MCF-7 |

**Supplemental Table S6: ENCODE histone ChIP-seq experiments in seven cell types** — See also Fig. S4. The ENCODE accession numbers allow to uniquely identify the experiment and peak call file (bigBed) on the ENCODE portal (https://www.encodeproject.org/).
